# Profiling plasma extracellular vesicle-derived microRNAs for noninvasive diagnosis of alpha-1 antitrypsin deficiency associated liver disease

**DOI:** 10.1101/2023.08.31.555751

**Authors:** Regina Oshins, Zhiguang Huo, Brittney Poole, Virginia Clark, Huiping Zhou, Jesse West, Matthew Wheeler, Mahya Aghaee, Mei He, Mark Brantly, Nazli Khodayari

## Abstract

Alpha-1 antitrypsin deficiency (AATD) is a rare genetic disease characterized by reduced circulating levels of alpha-1 antitrypsin (AAT) due to the retention of misfolded AAT in the hepatocytes. The toxic AAT aggregates in hepatocytes cause liver fibrosis, cirrhosis, and hepatocellular carcinoma. Most patients remain asymptomatic until the final stage in which liver transplantation becomes the only treatment. Timely diagnosis of AATD liver disease plays a critical role in intervention and finding curative solutions. Assessing the prevalence and severity of AATD liver disease remains challenging due to limitations in current methods. Liver biopsy, the gold standard for evaluating the hepatic AAT accumulation, the initiating stage for AATD liver disease, is hindered by invasiveness and sampling errors. To address these limitations, we conducted a study using unique and precious clinical samples. We analyzed plasma extracellular vesicle (EV) derived miRNAs and liver transcriptomes from AATD individuals to develop a sensitive and noninvasive diagnostic approach for AATD liver disease. In the testing stage of our study, we enrolled 17 AATD individuals with different stages of liver disease, as determined by liver biopsy, and 20 controls. We identified differential expression of 178 miRNAs within the AATD group compared to controls by miRNA sequencing. Among those miRNAs, we selected miR-223-3p, miR-23a-3p, miR-15b-5p, let-7a-5p, let-7f-5p, and miR-374a-5p for further validation in an independent cohort of 45 AATD individuals. Using a logistic model that combined three miRNAs, we achieved an AUC of 0.737 for detecting AATD liver disease. Adding a fourth miRNA into this model increased the AUC to 0.751. The changes in EV miRNAs are correlated to dysregulated expression of liver mRNAs in AATD individuals with different stages of liver disease. We propose that plasma-circulating EV exhibit distinct miRNAs in AATD individuals and could serve as clinically significant biomarkers for the early detection of AATD liver disease.

## INTRODUCTION

Alpha-1 antitrypsin deficiency (AATD) is a genetic disorder with a high prevalence in northwestern European countries and North America (1). AATD is caused by a single mutation of the SERPINA1 gene, resulting in misfolding and polymerizing of the Z mutant form of alpha-1 antitrypsin (AAT) in hepatocytes rather than appropriate secretion (2). While intrahepatic accumulation of misfolded AAT in AATD patients can lead to the development of liver disease, resultant low levels of AAT in the circulation leads to lung injury (3) (4). In adult AATD individuals, the induced liver disease is often underdiagnosed until the last stage of cirrhosis, which is untreatable other than by liver transplantation (5). In AATD children, some may present jaundice or develop liver cirrhosis, consequently requiring liver transplantation (6).

Liver fibrosis and its end-stage cirrhosis are major causes of morbidity and mortality, with limited therapeutic options, often leading to hepatocellular carcinoma (HCC) and increased mortality rates (7). While cirrhosis is an irreversible process, new evidence indicates that liver fibrosis is a dynamic and reversible process (8). Abnormal accumulation of AAT in hepatocytes has been identified as a cause of hepatocyte injury, leading to liver fibrosis in AATD patients (9). Therefore, it is crucial to assess the extent of hepatic AAT burden and identify AATD individuals in the early stage of liver disease to implement curative treatments before cirrhosis develops. Currently, liver biopsy is considered the “gold standard” for diagnosing AATD-mediated liver disease and assessing the grade of fibrosis (9). However, liver biopsy is an invasive surgical procedure associated with various complications, including abdominal pain, bleeding, and, rarely, death (10). Additionally, the limited size of liver samples obtained and the subjective assessment by pathologists limits the accuracy and reproducibility of histologic diagnosis (11). Given these challenges, alternative approaches are needed to overcome the limitations of liver biopsy in diagnosing and assessing fibrosis in AATD patients.

Extracellular vesicles (EV), which have diameters ranging from 30 to 200 nm, are released by all living cells (12). EV are known to selectively carry important signaling biomolecules such as microRNAs (miRNAs), mRNAs, proteins, and lipids. Most bio-fluids, including plasma and serum, contain a significant number of EV, which coordinate multiple physiological and pathological processes through transferring of their cargo between different cells and organs (4, 13). In the liver, almost all types of liver cells, including hepatocytes, both release and receive EV. EV have been shown to have an important role in the initiation and progression of liver diseases, including liver inflammatory disorders and liver fibrosis (13, 14). The study of EV contents has emerged as an area of interest in discovering new biomarkers in liver disease (13, 15–19).

It has been reported that different miRNAs are associated with EV, and their composition can vary across specific pathological conditions (20). Several EV-associated miRNAs have been found to be altered and linked to different types of liver diseases (10, 21). It is important to note that miRNAs, regardless of their etiology, have been associated with the clinical stage of various liver diseases, exhibiting diverse clinical symptoms, such as metabolic chronic liver disease and viral hepatitis (21). AATD-mediated liver disease is characterized by a toxic “gain-of-function” mechanism involving E.R. stress, oxidative stress, mitochondrial dysfunction, and lipotoxicity (22). However, there has been no systematic screening of plasma EV miRNA profiles specifically indicating the hepatotoxic features associated with AATD. Additionally, there is no clear consensus on the potential use of circulating miRNAs as diagnostic indicators for AATD-associated liver disease.

In this prospective observational study, we employed an innovative approach to compare the miRNA profiles of plasma EV from AATD individuals with different stages of liver disease to those of healthy controls. Our hypothesis was that EV-derived miRNAs could serve as effective biomarkers for diagnosing different stages of AATD-mediated liver disease and identifying individuals at risk of disease progression. It is important to note that our objective was not to identify miRNA biomarkers that could distinguish AATD individuals from healthy controls. Instead, our integrative approach involved analyzing both AATD individuals and healthy controls, followed by filtering steps to identify miRNAs that could provide insights into the liver disease status of AATD individuals and aid in detection of AATD individuals at risk of developing pathological liver disease. Furthermore, we leveraged liver biopsy samples from the same AATD individuals and performed RNA-sequencing analysis on liver tissues representing different stages of liver disease. By integrating these two data sets, we identified genes and pathways associated with the severity of AATD-mediated liver disease that were modulated by plasma EV-derived miRNAs. These findings may have functional implications in understanding the pathophysiology of AATD liver disease.

## Materials and methods

### Human subjects

The protocol for this study was approved by the Clinical Research Ethics Committee of the University of Florida (IRB202101148) and written informed consent was obtained from each subject in accordance with the Declaration of Helsinki Principles (23). AATD individuals (n=62 age range 35-70 years; mean 59 years; male: female ratio 27:30) were recruited by pulmonologists specialized in AATD in the Pulmonary Division of the University of Florida at Shands Hospital. Diagnosis of AATD individuals was established according to AAT genotyping, isoelectric focusing of the serum proteins, and AAT serum levels using nephelometry (Behring Diagnostics, Marburg, Germany) using in-house standards and controls (24). Percutaneous liver biopsy was performed in a well characterized cross section of adults with AATD as we previously reported (9). A group of age- and gender-matched healthy individuals (n = 20; age-range 37-79; mean 52 years; male: female 8:12) from the Alpha-1 DNA and Tissue Bank were also recruited. Detailed clinical data of all patients and healthy controls enrolled in this study are shown in Table 1.

**Table 1.** Demographics table of the study subjects.

|  | **MM (n=20)** | **ZZ (n=62)** |
| --- | --- | --- |
| **Age (years)** | 52 (37 – 79) | 57 (35 – 70) |
| **Sex (F%)** | 60 | 61 |
| **Augmentation** | 0 | 39 |
| **FEV1 (%)** | N/A | 64 |

### Sample collection and processing

Blood samples from all participants were obtained following the guidelines set by ISEV - International Society for Extracellular Vesicles (25). The collected blood was centrifuged at 2500 x g for 10 minutes at room temperature, and the resulting plasma was carefully separated, aliquoted, and stored at −80°C. Plasma aliquots were later defrosted on ice, quickly swirled through a 37°C water bath until clear and then centrifuged for 5 minutes at 18,000 x g at 4°C. The supernatant was then removed and passed through a 0.8μm filter. The filtrate was again passed through a 0.22μm filter before EV extraction for RNA isolation. 4 mL of plasma was used per sample.

### EV isolation

EV were isolated from cryopreserved plasma using differential centrifugation as previously described (4). Briefly, plasma was first centrifuged at 2000 x g for 30 minutes at 4°C. The resulting supernatant was then passed through a 0.8µm filter. The filtered samples were collected into ultracentrifuge tubes and subjected to ultracentrifugation in a Beckman Coulter Optima XE90 Ultracentrifuge with an SW-40Ti swinging rotor at 110,000 x g for 2 hours at 4°C. The EV pellets were resuspended in PBS, filtered through a 0.22μm filter (Millipore, Billerica, MA) and centrifuged at 110,000 x g for 70 minutes at 4°C. The EV were resuspended in 200μL of PBS and conserved at −80°C for later use.

### EV characterization

Purified EV fractions were characterized for their size, concentration, presence of specific EV markers, and morphology. To check EV morphology and size, isolated EV were subjected to transmission electron microscopy (TEM) and Nanoparticle Tracking Analysis (NTA), as reported earlier (4). EV were analyzed for the presence of CD81, CD63, and TSG101 as specific EV markers and absence of E.R. marker Calnexin by western blot assay.

### RNase and proteinase treatment of isolated plasma EV

Free RNA was eliminated from plasma by a combined treatment of 100µg/ml of Proteinase K and 10µg/ml of RNase A. First, isolated EV were treated with Proteinase K (Qiagen, Hilden, Germany) for 30 minutes at 37°C, followed by RNase A treatment (Qiagen, Hilden, Germany) for 15 minutes at 37°C.

### EV protein quantification

The protein concentration of EV enriched fractions was quantified by Pierce BCA Protein Assay Kit (Thermo Scientific, Waltham, MA) according to manufacturer instructions. 4μL of each standard and lysed EV enriched sample was pipetted into a 96-well plate with 200μL of reaction buffer and mixed thoroughly. Then the plate was covered and incubated at 37°C for 30 minutes. The absorbance at 562nm was measured with a spectrophotometer (SpectraMax, Molecular Devices) and a standard curve was used for the determination of the protein concentration of each EV sample.

### RNA isolation and RNA library preparation

Total RNA from AATD liver tissues was extracted using a RNeasy Plus Mini Kit (Qiagen, Germantown, Maryland). The quantity and quality of the extracted RNA were determined using an Agilent Bioanalyzer (Agilent Technologies, Santa Clara, CA). Total RNA was used for polyadenylation and synthesis of double-stranded cDNAs according to Illumina’s TruSeq RNA Sample Prep guidelines (San Diego, CA). Liver tissue library preparation and RNA sequencing were performed at the University of Florida Interdisciplinary Center for Biotechnology Research.

RNA from purified EV fractions was isolated using an exoRNeasy Maxi Kit (Qiagen, Germantown, MD) and the quantity and quality of the RNA were determined using the Agilent RNA 6000 Pico Kit and Small RNA Kit on the Agilent Bioanalyzer. Sequencing libraries were constructed using approximately 6 ng of total EV RNA as input material for the RNA sample preparations using the Small RNA Library Prep Set for Illumina kit (New England BioLabs) following manufacturer instructions. Each adapter-ligated RNA was mixed with ProtoScript II reverse transcriptase, RNase Inhibitor, and first-strand synthesis reaction buffer (NEB, cat. No. E7560S) and incubated for 60 minutes at 50°C. Index codes were then added to each reaction product for sample differentiation. Size selection was performed using a 6% polyacrylamide gel. For miRNA, the bands corresponding to ∼150 bp were isolated. The library quality was assessed with both the Agilent Bioanalyzer 2100 and Qubit and sequenced using the Illumina Miseq 1 × 150 cycles V3 kit at the University of Florida Interdisciplinary Center for Biotechnology Research.

### RNA-sequencing and bioinformatics

After removing adapters using the cutadapt software (26), the small RNA sequencing reads were aligned to the genome reference consortium human build 38 (GRCh38) using bowtie2 software (27) with “--very-sensitive-local” option local alignment, which does not require reads to align end-to-end. The miRNA expression count data were extracted using HTSeq software (28, 29) with the miRBase Sequence Database (release 22) (30) as the gene annotation reference. The resulting miRNA expression data were further normalized using the count per million reads method, followed by log2(x+1) transformation, where x is the expression value and 1 is the pseudo count. The quality of the liver tissue RNA-Seq data was evaluated using FastQC prior to further downstream analysis (FastQC. [(accessed on 26 April 2010)]. http://www.bioinformatics.babraham.ac.uk/projects/fastqc/). Low-quality sequences were trimmed, and poor-quality reads were removed using Trimmomatic (31). The Star Aligner was used to map high-quality paired end reads to the human genome of GRCh38 (32). Gene expression was obtained using RSEM (33).

### Functional Pathway, Upstream Regulator, and Network Analyses

We performed pathway enrichment analysis using Ingenuity Pathway Analysis (IPA) (Ingenuity Pathways Analysis System, Ingenuity Systems, Inc., Redwood, CA). The IPA core analyses are based on previous knowledge of the associations of upstream regulators and their downstream target genes archived in the Ingenuity knowledge base (34). An overlap p-value based on the significant overlap between the genes and their targets regulated by the transcriptional regulator and an activation z-score is computed. The activation z-score is used to infer likely activation or inhibition states of upstream transcriptional regulators. It was considered significantly activated or inhibited with an overlap p-value ≤ 0.05 and a z-score ≥ 2.0 (or ≤ −2.0). Using mRNA sequencing (RNA-seq) data, enrichment analyses were also conducted with the intent of exploring the functional pathways associated with various biological conditions. The specific genes of interest were identified in terms of statistical significance (adjusted p-value < 0.05) and substantial differential expression between groups. The differentially expressed genes (DEGs) obtained from the comparisons were then subjected to Kyoto Encyclopedia of Genes and Genomes (KEGG) pathway enrichment analysis using the enrich-KEGG (35) function from the R package cluster Profiler (36) with default parameters. Gene ontology (G.O.) enrichment analysis (based on biological process, cellular component, or molecular function) was conducted using hypergeometric distribution algorithms and the Fisher exact test. G.O. terms were assigned to each unique gene based on the G.O. annotation of the corresponding homologs in the database (37).

### Bibliographic search and miRNA selection

A literature search of the PubMed and Liver Disease MicroRNA Signature (38) databases was performed, aiming to identify miRNAs already described as deregulated in liver diseases. We then combined this bibliographic information with our results from NGS of plasma-derived EV from AATD and healthy control samples. Thus, a group of highly expressed miRNAs (all producing more than 5000 reads in the NGS assay and belonging to the topmost abundant miRNAs identified by NGS) were further considered for biomarker studies as an indicator for the severity of liver disease associated with AATD.

### Real-time qPCR validation of selected miRNAs

Reverse transcription reactions were performed using TaqMan Advanced miRNA Assays (Applied Biosystems Inc., CA, USA) using 2 μL of cell-free RNA extracted from EV to prepare cDNA. Real-time PCR reactions were performed in duplicate in 20 μL reaction volumes using 10 μL TaqMan Fast Advanced Master Mix, 1 μL TaqMan Advanced miRNA assay (20x) (Applied Biosystems. Inc., CA; USA), 4 μL of nuclease free water and 5 μL of R.T. product after a 1:10 dilution. Real-time PCR was carried out on an Applied BioSystems 7500 Fast thermocycler (Applied Biosystems. Inc., CA; USA) programmed as follows: 95°C for 20 s followed by 40 cycles of 95°C for 1 s and 60°C for 20 s. We used hsa-miR-21-5p and hsa-miR-26a-5p, two of the most stable miRNAs in terms of read counts, as endogenous controls. All fold-change data were obtained using the delta-delta C.T. method (2^-ΔΔCt^) 28. Five differentially expressed miRNAs detected after small RNA-seq were validated using RT-qPCR (4).

### Liver histology

A percutaneous liver biopsy was performed with a 16-gauge BioPince™ core biopsy needle. The sample was fixed in formalin and processed for examination. Stains included H&E, Masson’s trichrome, and PAS/PAS + D. Two pathologists (C.L., A.C.) scored each biopsy for fibrosis using METAVIR stages F0-F4. Clinically significant fibrosis was defined as stage ≥F2. PAS-positive, diastase-resistant (PAS + D) staining and immunohistochemistry (IHC) identified AAT accumulation. PAS + D globules present in a high-power field were scored from 0–3 as follows: 0-None, 1-Rare: <5 hepatocytes with globules, 2-Few: 5–20 hepatocytes with globules, 3-Numerous: ≥20 hepatocytes with globules (9).

### Statistical analysis

After removing non-expressed miRNAs, differential analysis was performed using the edgeR package (39), which employs a negative binomial to directly model the count data. Each miRNA’s predictive power was determined by a receiver operating characteristic (ROC) curve. To ensure an accurate and reliable ROC analysis, the miRNA data was normalized by CPM (count per million reads), log2 transformed, and standardized. In the ROC curve, the PASD status (no liver disease [PASD 0-1] and liver disease [PASD 2-3]) was the outcome variable, the expression level of a miRNA was the predictor. By varying the miRNA expression criteria in prediction PASD status, we could obtain a series of prediction sensitivities and specificities. The optimum sensitivity and the specificity was chosen such that the Youden index was maximized, where Youden index = sensitivity + specificity + 1. To assess the over predictive power, we further calculated the area under the curve (AUC). An AUC of 1 indicates perfect prediction, while an AUC of 0.5 is equivalent to random guessing. To explore the combined predictive power of miRNA panels (e.g., sets of 3 or 4 miRNAs), we constructed composite scores using logistic regression and assessed their performance using ROC analysis. To ensure the validity of our findings and avoid overfitting, we tested these composite scores on an independent sample of 20 AATD cases. For the RNASeq data, we similarly performed differential expression (DE) analysis using edgeR (39, 40). The thresholds were set at FDR <= 0.05 and a fold change of >= 2.

## RESULTS

### Isolation and characterization of EV

EV were isolated from the plasma of 20 control subjects (MM) and 62 AATD individuals (ZZ) with different stages of liver disease by a combination of filtration and ultracentrifugation and were characterized by analyses of particle size, distribution, and concentration using NTA as previously described (4). EV isolated from both MM and ZZ subject groups were within the normal range for EV size (below 200 nm) and the EV recovery from plasma in ZZ individuals was the same as those in the MM control group (Fig.1A). The presence of EV markers was investigated by western blot analysis. Results confirmed abundant CD63, CD81, and TSG101 expression in all EV fractions, while Calnexin (EV negative control) was not detected in EV fractions (Fig. 1B). To eliminate free miRNA and proteins that may be attached to EV, we performed a degradation assay using Proteinase K and RNase A. Coomassie blue staining showed that Proteinase K treatment degraded most proteins in plasma and isolated EV (Fig. 1C). After Proteinase K and RNase treatment, we also observed that control EV isolated from the plasma of the MM group had slightly higher concentrations of total RNA as compared to EV from the plasma of the ZZ group (Fig. 1D).

**Figure 1.**
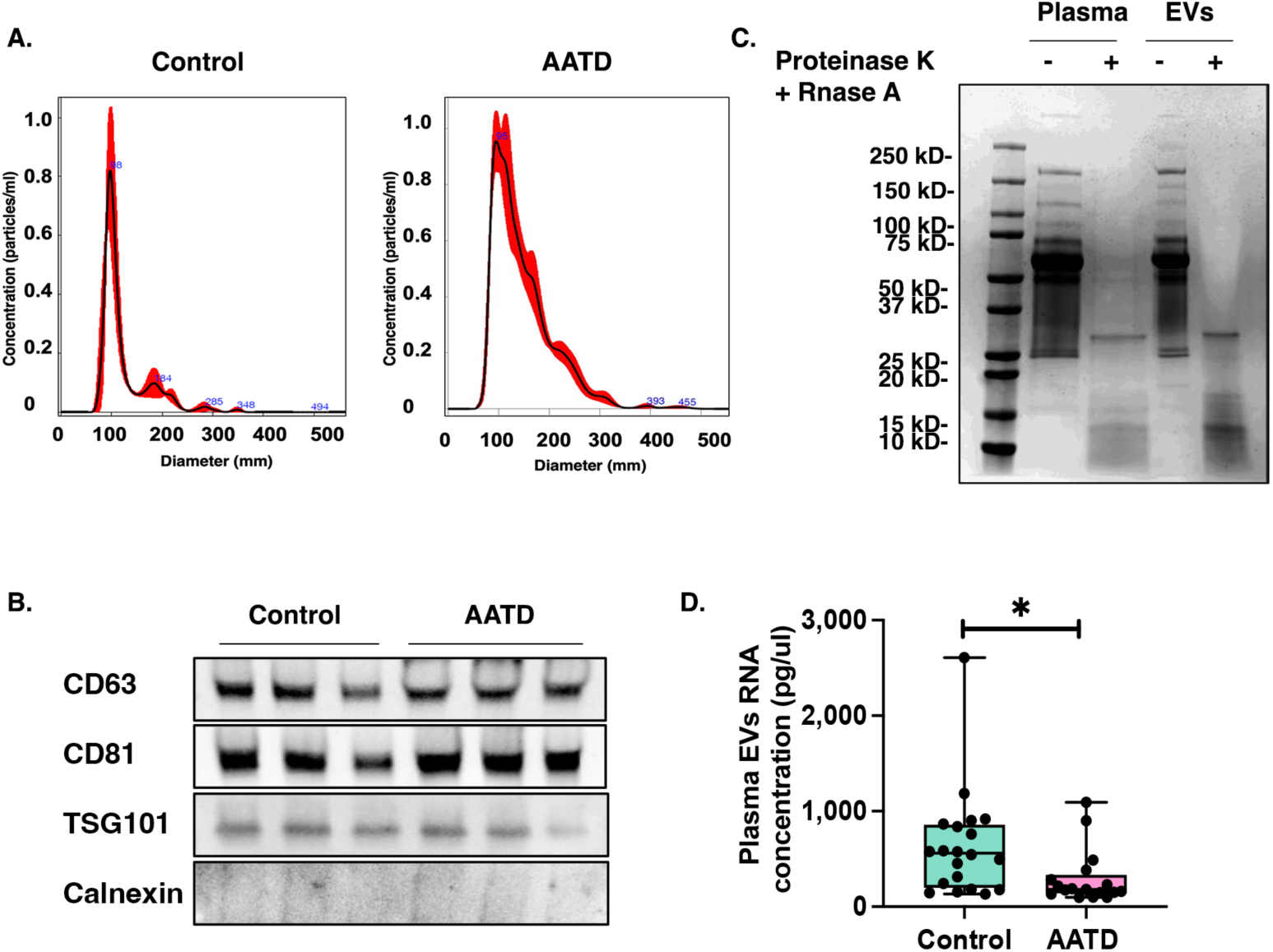
Isolated Extracellular Vesicles (EV) enriched fractions from participants’ plasma. **A.** Nano tracking analysis results suggested that EV isolated from plasma were about 75-200nm in diameter. **B.** EV surface markers CD63, CD81 and TSG101 were all detected by western blot in the EV fractions of the plasma, and Calnexin, a negative marker of EV was absent in our isolated EV fraction samples. **C.** Isolated EV fractions were treated with RNase A and protease K to eliminate EV’s membrane bound RNAs and proteins. **D.** A box plot comparing the RNA concentration (pg/µl) of EV fractions isolated from control and alpa-1 antitrypsin deficient (AATD) individuals.

### Plasma circulating EV associated miRNA profile of AATD and control individuals

To get a global profile of the plasma EV-derived miRNA from AATD individuals, we analyzed 37 samples (17 AATD, 20 controls) by small RNA sequencing. Based on the raw data, an average of 6,116,653 raw reads per sample were obtained for control samples and 11,846,649 reads per sample for the AATD group (Additional file 1: Table S1). A total of 682 mature known miRNAs were identified among all samples. More than 80% of these miRNAs (547 miRNAs) were already included in the vesicular database EVpedia as related to EV from human samples (Fig. 2A). Among the identified miRNAs, the hsa-let-7 family appeared as one of the most prevalent miRNA groups in our dataset, representing 48% of the miRNAs identified (Fig. 2B). This finding is in line with previous studies that have reported a correlation between plasma circulating levels of let-7 family and the severity of hepatic fibrosis (41). The complete list of identified miRNAs has been submitted to the EVpedia database under the title of this manuscript. In differential expression (DE) analysis (|log2(F.C.) |>1, p < 0.05), 178 miRNAs were found differentially expressed (Table 2), of which 92 miRNAs were up-regulated, and 86 miRNAs were down-regulated in AATD individuals compared to controls (Fig. 2C). Our small RNA-seq data indicated that miR-25-3p, miR-99b-3p, and miR-30a-5p are highly upregulated and miR-451a, miR-374a-5p, miR-146a-5p, miR-16-5p, miR-26b-5p, miR-96-5p, and miR-142-5p are highly downregulated in the plasma EV of AATD individuals as compared to healthy controls (Supplementary Fig.1A). The dysregulated expression of these miRNAs has been previously reported in various liver diseases (11, 42–54). To gain insights into the potential biological implications of the top dysregulated miRNAs, we performed core pathway analyses using Ingenuity Pathway Analysis (IPA) software. The association of dysregulated miRNAs with different liver disease pathophysiology is illustrated in Figure. 2D. All diseases associated with top dysregulated miRNAs in AATD individuals are illustrated in Supplementary Figure 1B.

**Figure 2.**
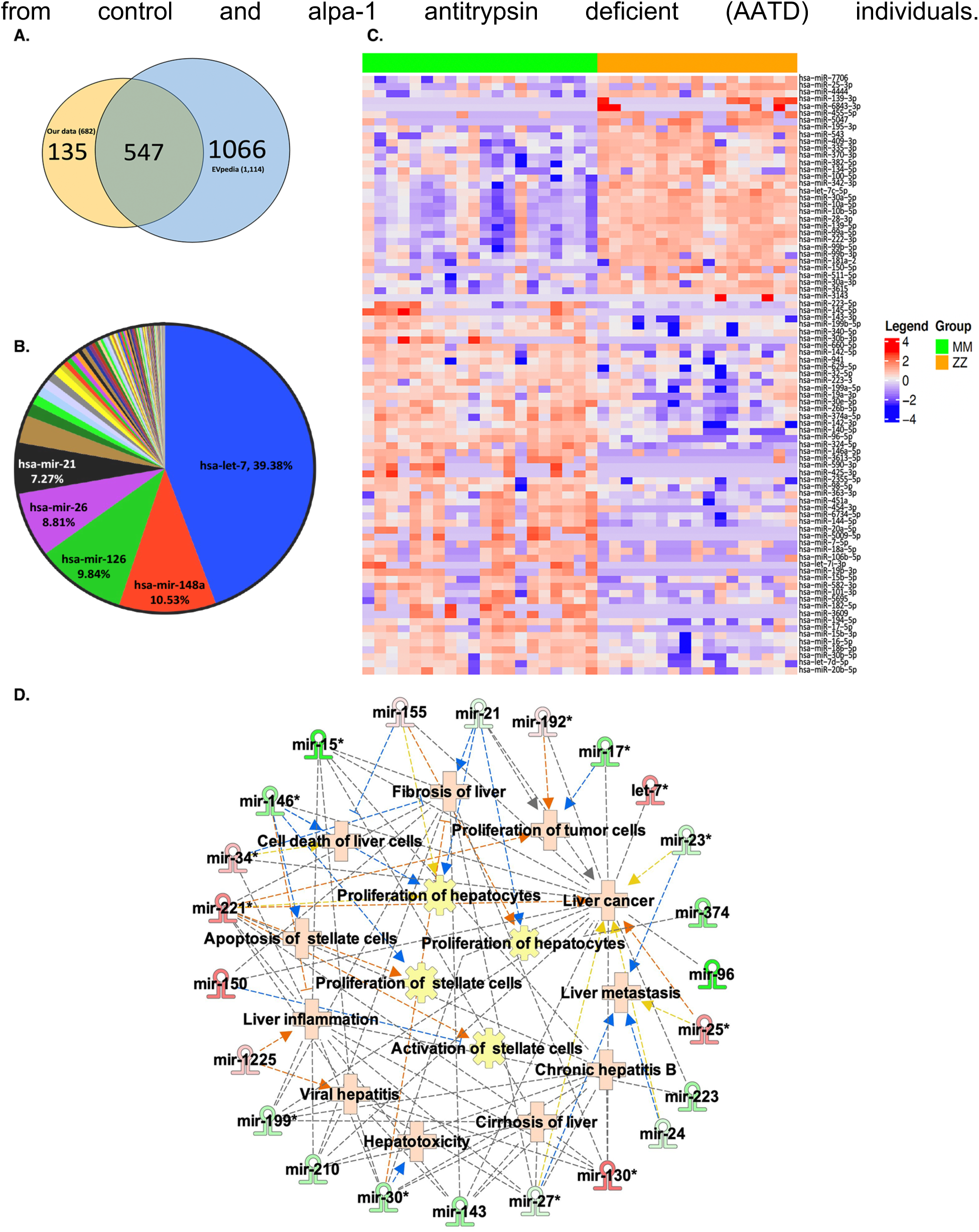
Plasma circulating Extracellular Vesicles (EV) associated miRNA profile of alpha-1 antitrypsin deficient (AATD) and control individuals. **A.** Venn diagram showed differentially expressed miRNAs (DEMs) shared between EVpedia dataset and our plasma EV fractions miRNA dataset. **B.** Percent abundance of the DEMs expression our plasma EV fractions miRNA dataset. **C.** Heatmap of the 100 DEMs expression level across 37 samples in our plasma EV fractions miRNA dataset. **D.** Pathway enrichment analysis from top 3 upregulated and top 7 downregulated DEMs across 37 samples in our plasma EV fractions miRNA dataset. Red indicates upregulated miRNAs and green indicates downregulated miRNAs from our dataset.

**Table 2.** Differential expression analysis (number of DEMs).

|  | **Total** | **Up regulated** | **Downregulated** |
| --- | --- | --- | --- |
| **q<0.05** | 85 | 31 | 54 |
| **q<0.1** | 120 | 54 | 66 |
| **q<0.2** | 173 | 90 | 83 |
| **p<0.001** | 57 | 342 | 37 |
| **p<0.01** | 102 | 342 | 61 |
| **p<0.05** | 179 | 342 | 84 |

### Validation of differentially expressed miRNAs by RT-qPCR

To validate our small RNA sequencing results, twenty miRNAs were randomly chosen. Some of these miRNAs have already been reported to be dysregulated in different types of liver diseases. Relative expression of these miRNAs was determined by RT-qPCR measurements using hsa-miR-26a-5p as the reference miRNA (55). All measurements were done in triplicate. Expression of hsa-miR-125a, hsa-miR-128-3p, hsa-miR-99a-5p, hsa-miR-335-5p, hsa-miR-361-3p, hsa-miR-150-5p, hsa-miR-30e-5p, hsa-miR-193a-5p, and hsa-miR-23a-3p were significantly upregulated in AATD individuals (Fig. 3A) compared control samples (Mann–Whitney p-values <0.001). Expression of hsa-miR-16-5p and hsa-miR-142-3p was significantly downregulated in AATD individuals (Fig. 3B) compared with control samples (Mann–Whitney p-values <0.001). Despite statistically significant differences in miRNA expression identified by small RNA sequencing, qPCR validation failed to replicate the difference of hsa-miR-96a-5p, hsa-miR-320a-3p, hsa-miR-146a-5p, hsa-miR-106b-5p, and hsa-miR-let-7d-5p between the two groups (Fig. 3C). However, 15/20 candidates show the same directional change in the RT-qPCR assay as identified in the miRNA-sequencing study.

**Figure 3.**
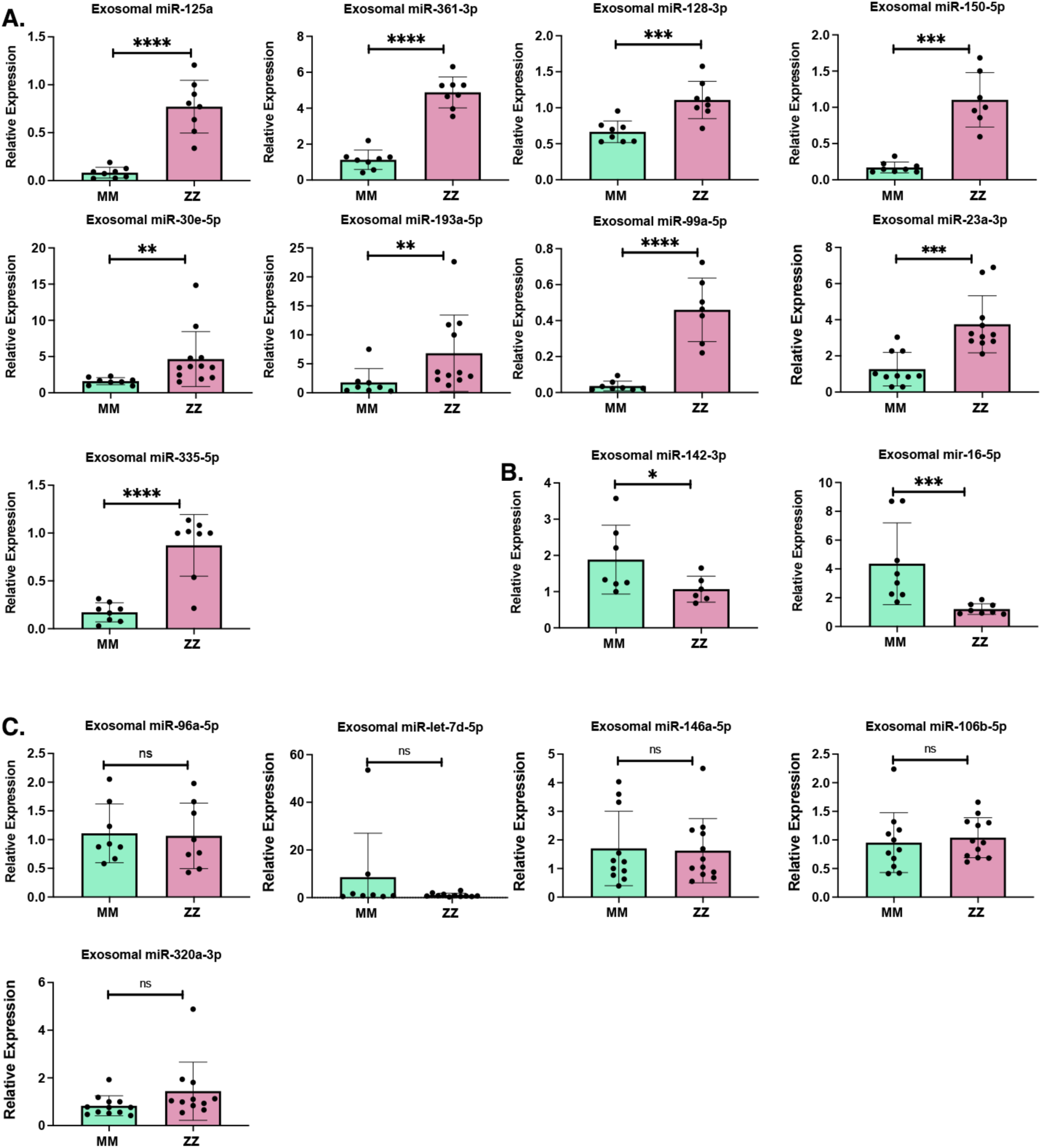
Validation of plasma Extracellular Vesicles (EV) fractions differentially expressed miRNAs by RT-qPCR. **A.** qPCR results from upregulated, **B.** Downregulated and **C.** Non-significant differential expressed miRNAs from our plasma EV fractions miRNA dataset.

### Target identification and functional enrichment

One hundred and eighty-seven miRNAs showing p<0.05 and log2 fold change >1 or <−1 (linear fold change >2 or <−2) were considered differentially expressed miRNAs (DEMs). The 178 DE miRNAs were further subjected to core pathway analyses using IPA software. We performed functional enrichment, target identification, and gene network analysis for the control and AATD plasma EV-derived miRNAs to further investigate molecular similarities/differences among the groups. Upstream regulators of these 178 DEMs are listed in additional file 2: Table S2. Causal networks for DEMs are listed in additional file 3: Table S3. Top significant canonical pathways for upregulated and downregulated miRNAs in control and AATD plasma EV are illustrated in Fig. 4A. Significantly altered pathways included the HOTAIR regulatory pathway and necroptosis signaling pathways. Additionally, we analyzed the potential impact of up-regulated and down-regulated miRNAs in AATD individuals to determine the potential function of those miRNAs on protein-coding mRNAs. IPA revealed that AATD plasma circulating EV-associated miRNA signatures are associated with the target genes linked with cellular and organismal functions, such as cell death and survival, cellular development, growth and proliferation, molecular transport, protein degradation and synthesis, lipid metabolism, and infectious diseases (Fig. 4B). Dysregulated miRNAs in AATD plasma circulating EV were also associated with liver, cardiac, and renal dysfunction as indicated by the tox function feature of IPA software (Fig. 4C).

**Figure 4.**
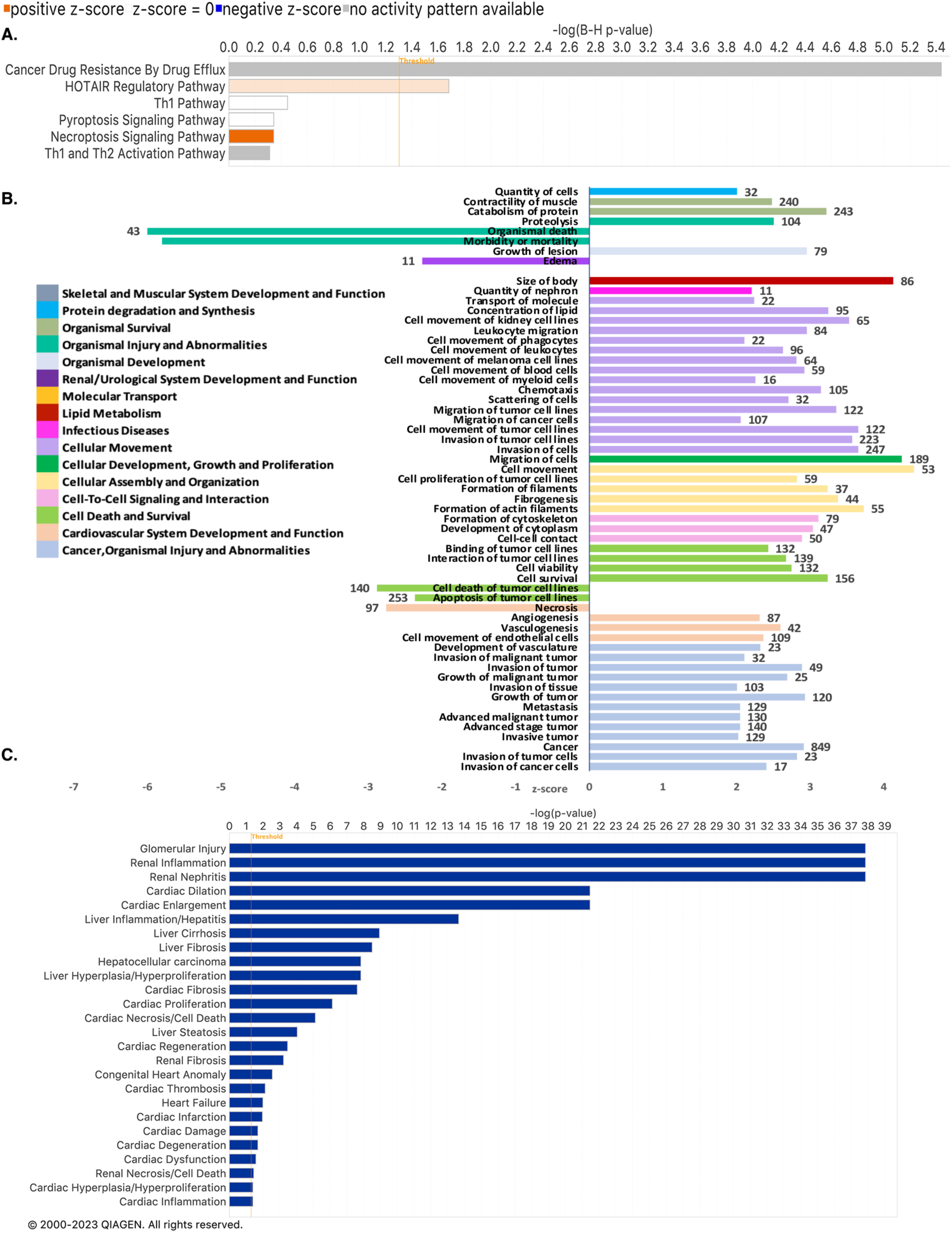
Pathway identification and functional enrichment. **A.** Canonical pathways targeted by the 178 differentially expressed miRNAs (DEMs) identified in our plasma extracellular vesicles (EV) fractions of alpha-1 antitrypsin deficient (AATD) individuals as predicted by Ingenuity Pathway Analysis (IPA). **B.** Bar plot of gene ontology (Cellular Component and Molecular Function) enriched. **C.** A bar plot shows the numbers of diseases and abnormalities associated with DEMs in our plasma EV fractions miRNA dataset.

### Correlation of plasma EV miRNA with PASD and fibrosis scores in AATD individuals

As we previously have shown (9), AATD-mediated liver disease is characterized by the accumulation of misfolded AAT in the hepatocytes, forming PASD (Periodic Acid-Schiff Diastase) globules and subsequent liver fibrosis (Fig. 5A). We analyzed the associations between the differentially expressed EV-associated miRNAs and PASD and fibrosis scores in AATD individuals. We identified 89 miRNAs associated with PASD scores and 68 miRNAs associated with fibrosis scores in AATD individuals. Notably, there was an overlap of 60 miRNA that were associated with both PASD and fibrosis scores (Fig. 5B). The top 10 miRNAs associated with PASD scores are shown in Table 3. The top 10 miRNAs associated with fibrosis scores are shown in Table 4. IPA target analysis revealed that these 60 miRNAs associated with both PASD, and fibrosis score predominantly target genes involved in proliferation and fibrogenesis (Fig. 5C).

**Figure 5.**
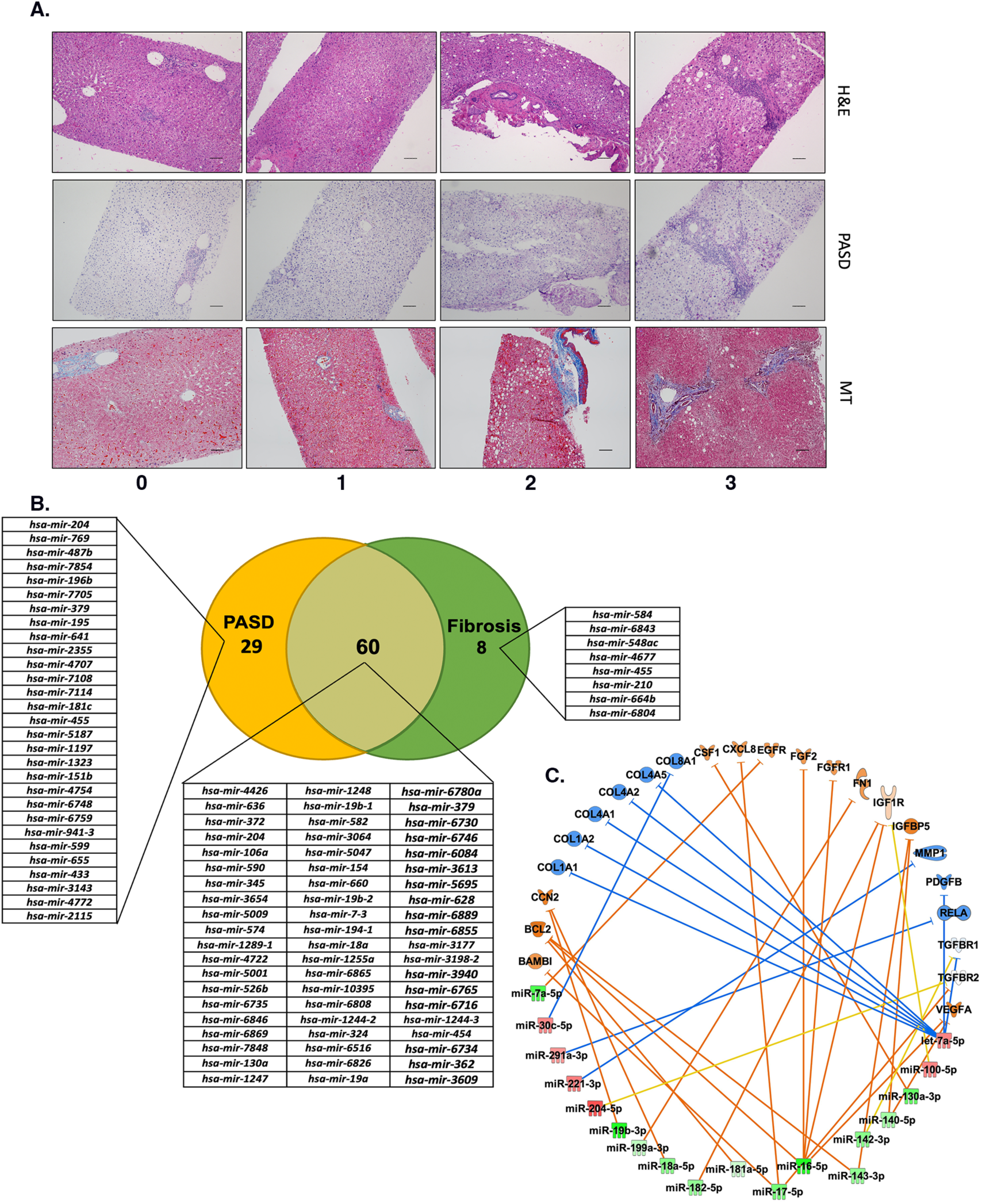
Correlation of plasma Extracellular Vesicles (EV) miRNA with PASD and fibrosis scores in alpha-1 antitrypsin deficient (AATD) individuals. **A.** Representative liver biopsies show increasing fibrosis stage and PASD globule score from AATD individuals. Top (Hematoxylin & Eosin), middle (PAS-D), bottom (Masson’s Trichome). Scale bar represents 200 µm. **B.** The associations between the differentially expressed EV-associated miRNAs and PASD, and fibrosis scores in AATD individuals. **C.** Ingenuity target analysis indicating target genes that are involved in proliferation and fibrogenesis. Red indicates upregulated miRNAs and green indicates downregulated miRNAs from our dataset. Orange color indicates upregulation and blue indicates downregulation of target genes, predicted by Ingenuity Pathway Analysis (IPA) software.

**Table 3.** Top 10 significant miRNA of EV derived from plasma of AATD individuals associated with the PASD score.

| miRNA_ID | Fold Change | P value |
| --- | --- | --- |
| hsa-miR-1248 | -6.5395301 | 3.68E-06 |
| hsa-miR-19b-1 | -3.6330913 | 6.40E-05 |
| hsa-miR-19b-2 | -3.6484493 | 0.000169582 |
| hsa-miR-582 | -2.7648581 | 0.000249953 |
| hsa-miR-660 | -2.213101 | 0.000371254 |
| hsa-miR-7-3 | -2.7947088 | 0.000779255 |
| hsa-miR-3064 | 1.52922162 | 0.000780172 |
| hsa-miR-5047 | 1.79885521 | 0.0009429 |
| hsa-miR-19a | -1.5003409 | 0.00100871 |
| hsa-miR-454 | -1.8302099 | 0.001019201 |

**Table 4.**
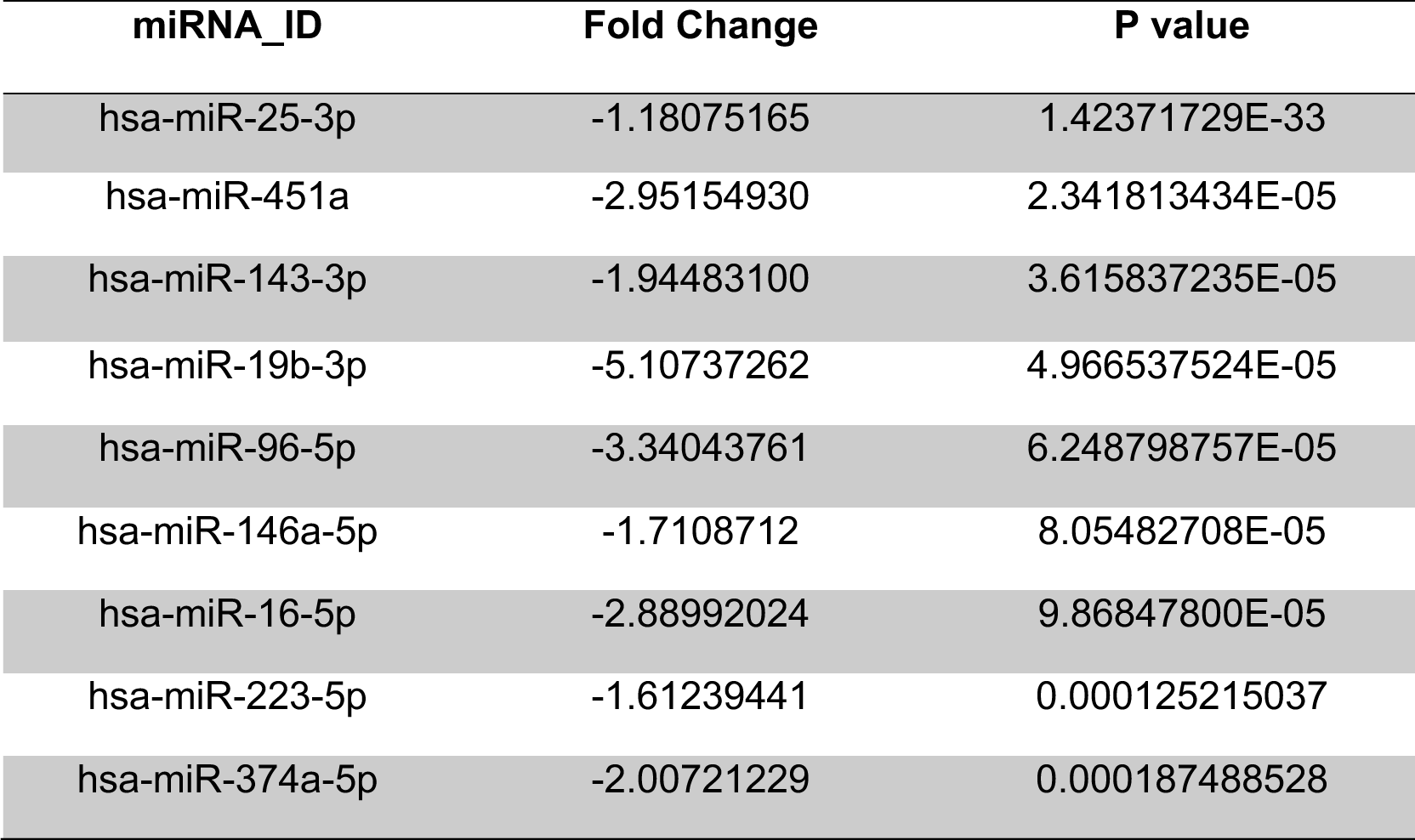
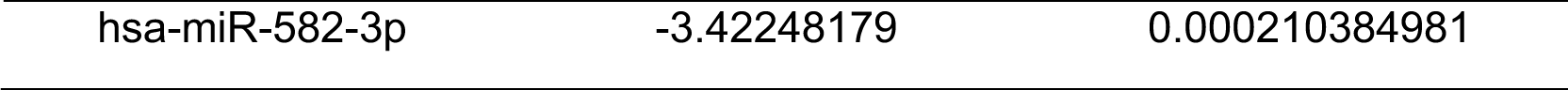
Top 10 miRNA of EV derived from plasma of AATD individuals associated with the fibrosis score.

### Identification of plasma EV-associated miRNA biomarker candidates for AATD liver disease severity

We first divided the experimental set of 17 AATD individuals into two groups: no liver disease (PASD 0-1) and liver disease (PASD 2-3). In DE analysis (|log2(F.C.) |>1, p < 0.05), 39 miRNAs were found to be differentially expressed (Fig. 6A). Next, for the screening of potential biomarkers in AATD-mediated liver disease diagnosis, we calculated the specificity and sensitivity of each DEM. Most candidate miRNAs displayed a specificity of 0.7–1.0 and a sensitivity of 0.8–1.0. To verify the potential of miRNA expression, ROC analysis was performed, and AUC was calculated (Table 5). A total of 6 miRNA were selected based on their significant differential expression and theoretical evidence for clinical application in previous studies (56) (Table 6). To validate the specificity of these 6 plasma EV-derived miRNAs for AATD individuals found in the testing set, ROC analysis was performed, and AUC was calculated (Fig. 6B) in both the testing and validation batches (n=62). As a next step to develop a better classification method, we used Fisher’s linear discriminant analysis to design comprehensive classifiers consisting of 3 and 4 miRNAs in the training set (Additional files 4 and 5: Tables S4 and S5) and the best results of the three miRNA combinations are shown in Figure 6C and the four miRNA combinations in Figure 6D.

**Figure 6.**
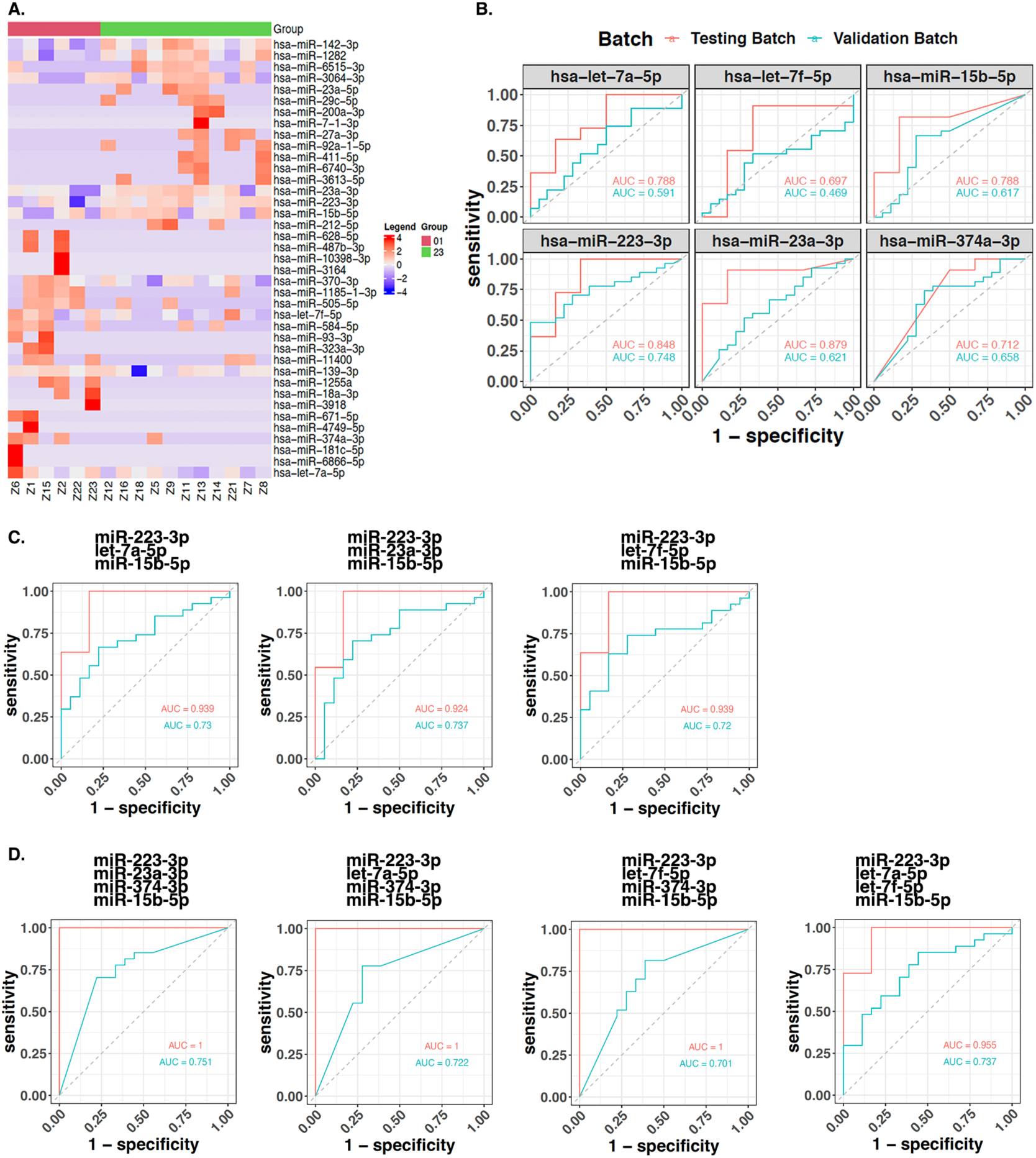
Identification of plasma Extracellular Vesicles (EV)-associated miRNA biomarker candidates for alpha-1 antitrypsin deficient (AATD) liver disease severity. **A.** A heatmap of the 36 differentially expressed miRNAs (DEMs) expression levels across 45 samples of the validation set in our plasma EV fractions miRNA dataset. **B.** Verification of EV derived hsa-let-7a-5p (A), hsa-let-7f-5p (B), hsa-miR15b-5p (C), hsa-miR223-3p (D), hsa-miR23a-3p (E), and hsa-miR374a-3p (F) as biomarkers of AATD- mediated liver disease. The ROC curves of the testing dataset (miRNA sequencing of 17 samples) were shown in red, while the ROC curves of validation dataset (miRNA sequencing of 45 samples) were shown in turquoise. **C.** Verification of 3 miRNAs panel 1 (miR223-3p + let-7a-5p + miR15b-5p), panel 2 (miR223-3p + miR32a-3p + miR15b-5p), and panel 3 (miR23a-3p + let-7f-5p + miR15b-5p) as biomarkers of AATD-mediated liver disease. **D.** Verification of 4 miRNAs panel 1 (miR223-3p + miR23a-3p + miR374-3p + miR15b-5p), panel 2 (miR223-3p + let-7a-5p + miR374-3p + miR15b-5p), panel 3 (miR223-3p + let-7f-5p + miR374-3p + miR15b-5p), and panel 4 (miR223-3p + let-7a-5p + let-7f-5p, miR15b-5p) as biomarkers of AATD-mediated liver disease. The ROC curves of testing dataset (miRNA sequencing of 17 samples) were shown in red, while the ROC curves of validation dataset (miRNA sequencing of 45 samples) were shown in turquoise.

**Table 5.** Top 24 miRNAs ranked by Sensitivity × Specificity (miRNA-seq data).

| miRNA_ID | log2FC | p value | Sensitivity | Specificity | Sen × Spe |
| --- | --- | --- | --- | --- | --- |
| hsa-miR-30a-5p | 1.73429 | 5.0516893E-06 | 1 | 0.9 | 0.9 |
| hsa-miR-10b-5p | 2.42141 | 0.00034663 | 1 | 0.9 | 0.9 |
| hsa-miR-361-3p | 1.1718 | 0.00768856 | 0.882352941 | 1 | 0.882352941 |
| hsa-miR-10a-5p | 2.16345 | 3.1185616E-05 | 1 | 0.85 | 0.85 |
| hsa-miR-186-5p | -2.1933 | 0.000117234 | 1 | 0.85 | 0.85 |
| hsa-let-7c-5p | 1.79436 | 0.000356463 | 1 | 0.85 | 0.85 |
| hsa-miR-451a | -3.94075 | 2.0757460E-07 | 0.941176471 | 0.9 | 0.847058824 |
| hsa-miR-222-3p | 1.89706 | 3.4847979E-05 | 0.941176471 | 0.9 | 0.847058824 |
| hsa-miR-3615 | 2.48389 | 5.9946212E-05 | 0.941176471 | 0.9 | 0.847058824 |
| hsa-miR-20a-5p | -2.28294 | 6.6973813E-05 | 0.941176471 | 0.9 | 0.847058824 |
| hsa-miR-146a-5p | -1.92759 | 1.5714056E-06 | 1 | 0.8 | 0.8 |
| hsa-miR-96-5p | -4.03593 | 2.5008768E-06 | 0.941176471 | 0.85 | 0.8 |
| hsa-miR-30a-3p | 2.0446 | 1.4232670535E-05 | 0.941176471 | 0.85 | 0.8 |
| hsa-miR-28-3p | 1.60626 | 0.000104222 | 1 | 0.8 | 0.8 |
| hsa-miR-125a-5p | 1.99188 | 0.006954756 | 0.941176471 | 0.85 | 0.8 |
| hsa-miR-99b-5p | 1.42507 | 0.000576658 | 0.823529412 | 0.95 | 0.782352941 |
| hsa-miR-99a-5p | 1.77386 | 0.000865078 | 0.941176471 | 0.8 | 0.752941176 |
| hsa-miR-1244 | 1.45254 | 0.005741546 | 0.941176471 | 0.8 | 0.752941176 |
| hsa-miR-16-5p | -3.75298 | 2.0737678880E-06 | 0.882352941 | 0.85 | 0.75 |
| hsa-miR-194-5p | -2.49102 | 4.8300450714E-05 | 0.882352941 | 0.85 | 0.75 |
| hsa-miR-370-3p | 1.87839 | 0.0007734 | 0.882352941 | 0.85 | 0.75 |
| hsa-miR-150-5p | 2.03927 | 0.001558803 | 0.882352941 | 0.85 | 0.75 |
| hsa-miR-139-5p | 1.57692 | 0.001821215 | 1 | 0.75 | 0.75 |
| hsa-miR-93-5p | -1.58784 | 0.006132461 | 1 | 0.75 | 0.75 |

**Table 6.** Circulating plasma miRNAs related to liver diseases.

| **miRNA_ID** | **Reported in patients' plasma (PMID)** | **Reported in liver disease (PMID)** |
| --- | --- | --- |
| hsa-let-7a-5p | Chronic Hepatitis C (30443558) | Hepatic Fibrosis (27227815) |
| hsa-let-7f-5p | Hepatocellular Carcinoma (32102538) | Hepatocellular Carcinoma (33303811) |
| hsa-miR-15-5p | Hepatocellular Carcinoma (26119771) | Liver Cancer (31814849) |
| hsa-miR-233-3p | Acute-on-chronic liver failure<br>(31919459) | Hepatocellular Carcinoma (29207133) |
| hsa-miR-23a-3p | NAFLD (27493762) | NAFLD (34723832) |
| hsa-miR-374a-3p | HBV-Liver fibrosis (30772950) | Hepatitis B-related acute-on-chronic<br>liver failure (26267843) |

### Liver transcriptomics of AATD individuals with different stages of liver disease

To assess the expression of the genes that associate with PASD score in AATD livers, we analyzed 50 AATD individuals’ liver samples by RNA sequencing. Our analysis showed that interferon alpha inducible protein 6 (IFI6) was the only differentially expressed gene when livers with PASD 1 were compared to those with PASD 0. Furthermore, there were 267 differentially expressed genes when livers with PASD 2 were compared to PASD 0, and 55 differentially expressed genes when comparing livers with PASD 3 to those with PASD 0 (Additional files 6: Table S6). We also found 36 common differentially expressed genes when livers with PASD 2 and 3 were compared to livers with PASD 0. Of all differentially expressed genes, 273 upregulated genes were associated with PASD. There were also 12 genes that were found to be downregulated in association with PASD scores (Fig. 7A). RNA-sequencing data was imported into Ingenuity Pathway Analysis which predicted activation of hepatic stellate cells in the liver of AATD individuals while liver disease is progressing (Supplementary Fig. 2A). We also ran pathway analysis for those 267 differentially expressed genes in the liver of AATD individuals with PASD 2 compared to those livers with PASD 0. Canonical pathways related to these genes have been illustrated in Supplementary Fig. 2B. Canonical pathways related to the 55 differentially expressed genes when comparing livers with PASD 3 to those with PASD 0 are illustrated in Supplementary Fig. 2C. To further enhance our understanding of the DEGs associated with our RNA-seq data set and their functional environment, we employed KEGG pathway enrichment analysis using the DEGs from the comparison of PASD 1 to PASD 2 liver tissues and PASD 2 to PASD 3 liver tissues. We observed that 10 pathways with p≤0.05 were significantly enriched in each comparison, including motor proteins, glycolysis/gluconeogenesis, AMPK signaling pathway, fructose, and mannose metabolism, and so on (Fig. 7B). The top 20 enriched terms were mainly involved in biological processes (muscle tissue development, myofibril assembly, and regulation of calcium ion transmembrane transport), cellular components (myofibril, stress fibers and myofilament) and molecular functions (extracellular matrix binding, chaperone binding, and lipoprotein receptor activity) (Fig. 7C).

**Figure 7.**
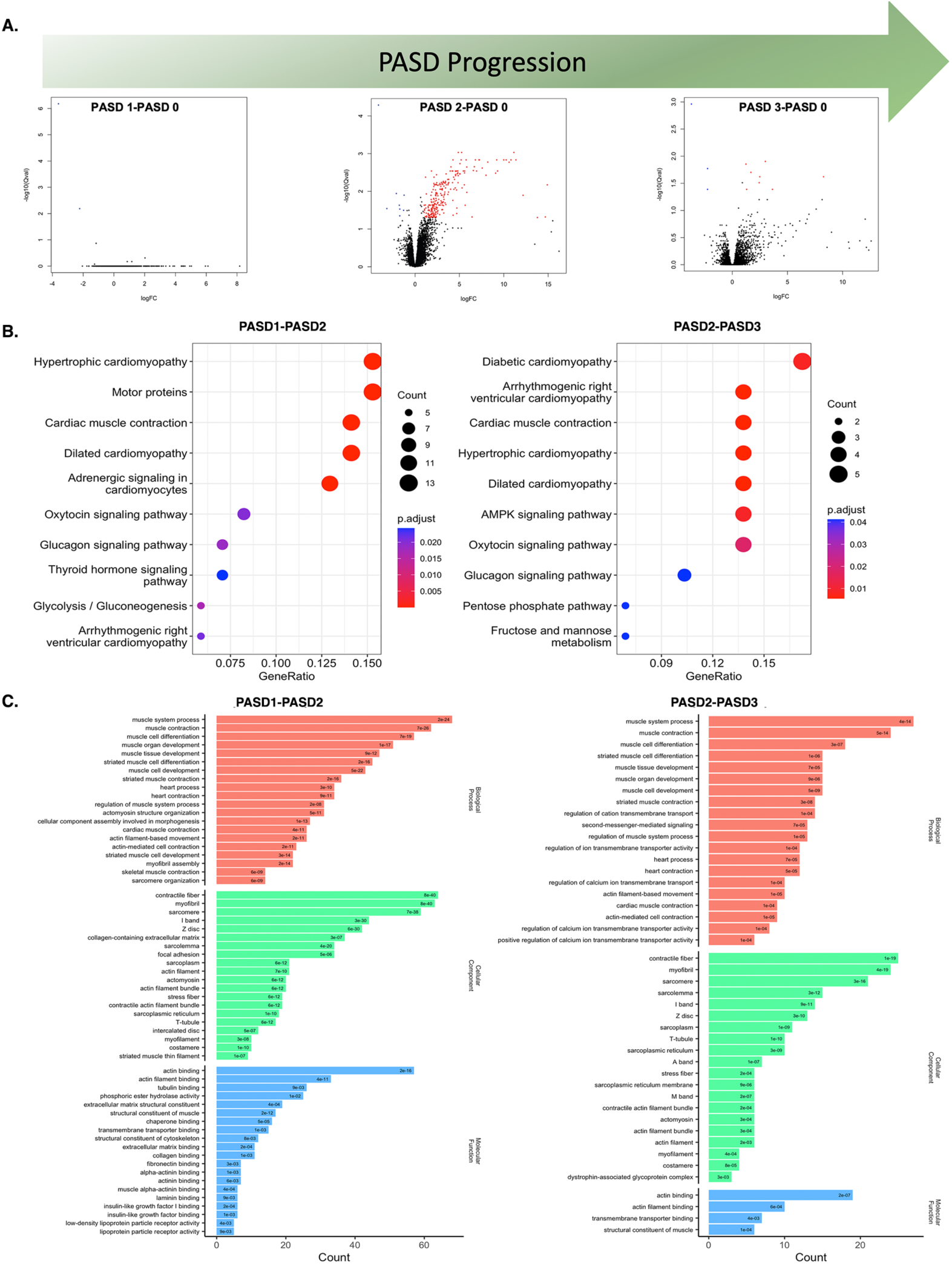
Correlation of liver transcriptomics with PASD in alpha-1 antitrypsin deficient (AATD) liver tissues. **A.** Changes in the expression of the genes in the AATD liver tissue samples during the progression of liver disease associated with AATD. **B.** KEGG pathway enrichment analysis using the differentially expressed genes (DEGs) from the comparison of PASD1 to PASD2 liver tissues and PASD2 to PASD3 liver tissues. **C.** Biological processes, cellular components, and molecular functions enriched using the differentially expressed genes (DEGs) from the comparison of PASD1 to PASD2 liver tissues and PASD2 to PASD3 liver tissues.

### The integration analysis of RNA-seq and miRNA-seq in AATD individuals

By integrating DEGs obtained from liver RNA sequencing into the circulating EV-associated DEMs, we found that multiple DEGs serve as targets for plasma EV-derived DEMs in AATD individuals. One hundred and eighty-seven DEMs and 55 DEGs identified in the previous steps were considered for the miRNA-mRNA integrated analysis. A total of 10734 genes were consistently predicted as potential targets of 39 DEMs using an online bioinformatics database miRWalk. Then a Venn diagram showed that 1190 overlapped genes were obtained between the predicted target genes and the identified DEGs (Fig. 8A), which were selected for further correlation analysis. According to the standard of absolute Pearson Correlation Coefficient (PCC) ≥ 0.8 and p-value ≤ 0.05, we finally identified 206 significant miRNA-mRNA pairs, including 53 negative correlating pairs and 153 positive correlating pairs (Fig. 8B). Since miRNAs generally suppress the expression of target mRNAs, those pairs where the miRNA and mRNA expression levels changed in the same direction were filtered out. In the case of 53 negative miRNA-mRNA interaction pairs, 21 pairs that composed 8 miRNA and 21 mRNA were up-regulated miRNAs vs. down-regulated mRNAs, while 32 pairs composed 16 miRNA and 27 mRNA were down-regulated miRNAs vs. up-regulated mRNAs (Additional files 7 and 8: Tables S7 and S8). Figure. 8C shows negative correlated mRNA-miRNA pairs from our AATD cohort that are involve in hepatic fibrosis. Remarkably, IPA also revealed that distinct pathways related to AATD liver disease including liver fibrosis, cirrhosis, and mitochondrial dysfunction might interact with each other (Fig. 8D).

**Figure 8.**
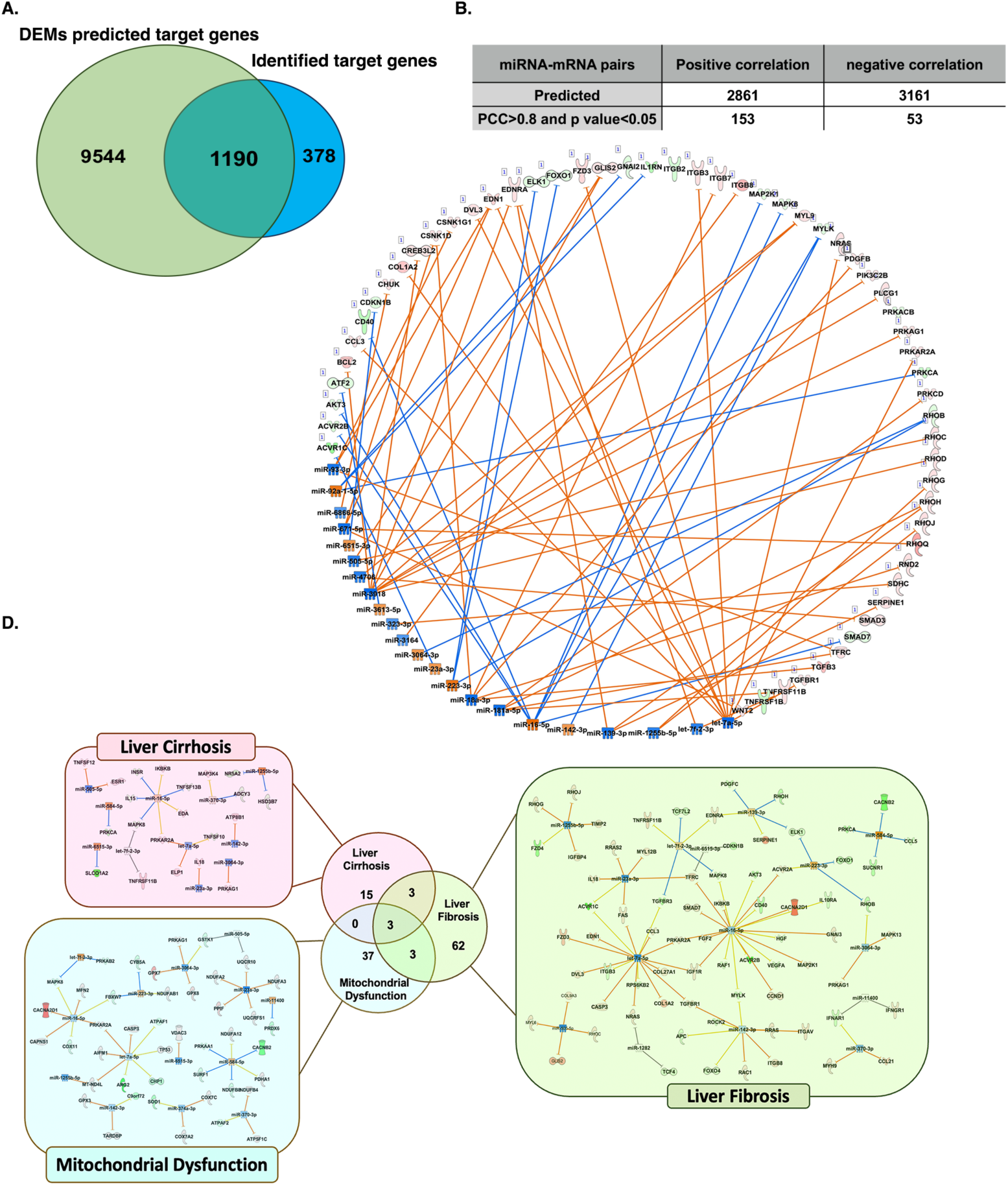
Integrated analysis of hepatic mRNA and plasma Extracellular Vesicles (EV)-associated miRNA expression profiles in alpha-1 antitrypsin deficient (AATD) individuals. **A.** Venn diagram between identified differentially expressed genes (DEGs) and target genes of differentially expressed miRNAs (DEMs). **B.** Pearson Correlation Coefficient (PCC) analysis to identify the negative correlating miRNA-mRNA pairs. Ingenuity Pathway Analysis (IPA) shows the interconnection of DEMs and DEGs in AATD individuals without liver disease (PASD 0 and 1) compared to with liver disease (PASD 2 and 3) **D.** Schematic overview of the miRNA-mRNA interaction networks in liver specific pathways. Red indicates upregulated miRNAs and green indicates downregulated miRNAs from our dataset. Orange color indicates upregulation and blue indicates downregulation of target genes, predicted by Ingenuity Pathway Analysis (IPA) software.

## DISCUSSION

In the present study, we isolated circulating EV from the plasma of AATD individuals and used the nomenclature “Generic term extracellular vesicle (EV),” according to the MISEV2018 guidelines (57) instead of the commonly used term “exosome” found in most related biomarker studies. Recently, EV have attracted much attention as a new noninvasive diagnostic tool to identify different asymptomatic diseases (15). The cargo of plasma circulating EV includes different types of nucleic acids and proteins. EV-associated miRNA has been the focus of biomarker-related studies because of its specificity, abundance, and stability (58). In this study, we conducted a comprehensive analysis of plasma circulating EV-associated miRNA profiles in individuals with AATD, comparing them to healthy controls. We specifically focused on selecting miRNAs that were differentially expressed in the plasma EV of AATD individuals with and without liver disease. These differentially expressed miRNAs were considered potential biomarker candidates for the detection of liver disease within the AATD population. We propose that biomarker panels of 3 and 4 miRNAs could serve as promising biomarkers for AATD-mediated liver disease. Furthermore, we analyzed the mRNA transcriptomics of the liver tissue from AATD individuals with and without liver disease. This analysis allowed us to explore the molecular pathways involved in the progression of liver disease in AATD individuals. By integrating the liver mRNA transcriptomics data with the plasma EV-derived miRNA data, we were able to identify the dysregulated miRNA-mRNA networks in the liver of AATD individuals. This comprehensive approach provides a deeper understanding of the molecular mechanisms underlying liver disease progression in AATD.

Next generation small RNA sequencing revealed a total of 178 miRNAs that are significantly dysregulated in the plasma EVs of AATD individuals compared to healthy individuals, suggesting that AATD status affects EV-mediated signaling in the plasma. Of DE miRNAs in the cohort of AATD individuals, we found that levels of EV-associated miR- 25-3p, miR-99-3p, and miR-30a-5p were significantly elevated. Most significant decreases were observed in EV-associated miR-451a, miR-374a-5p, miR-96-5p, and miR-16-5p. Of the top ten dysregulated miRNAs, miR-25-3p, miR-96-5p, and miR-30a- 5p have been reported as biomarkers for discriminating hepatocellular carcinoma patients (11, 42–51). Amongst them, miR-25-3p has been shown to promote hepatocellular carcinoma cell growth, migration, and invasion (54). Our data demonstrate the association of top dysregulated miRNAs in the plasma-derived EV from AATD individuals with hepatocellular carcinoma. This result is consistent with previous literature indicating that AATD-mediated liver disease is a risk factor for HCC (59). This also suggests the potential of plasma-derived EV-associated miRNA as an early marker to detect AATD individuals with a higher risk of HCC. MiRNA cargo of the plasma EV has been shown to represent disease-associated changes in their origin cells (60). Studies have shown that high extracellular level of miR-99-3p is linked to pericellular fibrosis in NAFLD (52), and low level of miR-16 is associated with the expression of hepatocyte growth factors in hepatic fibrosis (53). Given the fact that liver fibrosis is found in 35% of AATD individuals (9), dysregulation of EV-associated miR-99-3p and miR-16 in AATD plasma EV may represent a novel compelling liver fibrosis diagnostic option in these patients. Consistent with these results, pathway analysis from those 178 dysregulated miRNAs in the plasma EVs of AATD individuals also indicated that metabolic pathways related to cell migration, cell proliferation, and fibrogenesis were strongly upregulated, suggesting that miRNAs may be functional, playing a pivotal role in the regulation of hepatic fibrogenesis in AATD individuals (61).

Network analysis of the plasma EV-associated DEMs from AATD individuals suggested an increase in the activity of different canonical pathways in our data set, including the HOTAIR canonical pathway. Hox transcript antisense RNA (HOTAIR) is a long noncoding RNA implicated in gene silencing through binding to enzymes associated with chromatin remodeling (62). HOTAIR expression has been shown to be significantly increased in mouse models of liver fibrosis and human fibrotic livers and is associated with activated hepatic stellate cells (63). The mechanistic network of upstream regulators also demonstrated migration and activation of hepatic stellate cells, the initiation step in liver fibrogenesis. Our findings provide evidence indicating that the dysregulation of metabolic pathways and HOTAIR, are consistent with our previous observation that over one-third of AATD individuals have significant underlying liver fibrosis (9).

AATD patients with positive liver PASD scores and liver fibrosis show an important liver pathological overlap, hampering correct diagnosis, treatment, and clinical management (64). Although understanding the burden of AATD liver disease is important, fibrosis is also a critical issue to be addressed. We have previously reported that liver fibrosis is found in over one-third of AATD individuals and the presence of accumulated AAT in hepatocytes as PASD globules is associated with clinically significant fibrosis (9). To investigate this further, we analyzed the relationship between the circulating EV miRNA profile of AATD individuals and their PASD and fibrosis scores obtained from liver biopsy samples. Among analyzed plasma EV miRNAs, we identified 89 miRNAs significantly correlated with PASD score and 68 miRNAs significantly correlated with fibrosis score within the AATD cohort. Interestingly, we found 60 overlapping miRNAs associated with both PASD and fibrosis scores, further supporting our previous findings linking PASD globules and liver fibrosis in AATD individuals (9). Among these overlapping miRNAs, we observed upregulated expression of let-7a-5p. Intriguingly, let-7a-5p was predicted by IPA to be negatively and significantly correlated to collagen expression which serves as an indicator of fibrogenesis (Fig. 5C). Moreover, previous studies have shown that let-7a inhibits liver fibrogenesis through the TGFβ/SMAD signaling pathway (65). This suggests that let-7a-5p upregulation can be an anti-fibrotic defense mechanism in the liver of AATD patients. The differential expression of miRNAs in patients with different PASD and fibrosis scores may not only be a cause or specific consequence of each condition, but also a result of a common injury process within the liver of AATD individuals. It is known that different miRNAs can perform the similar functions or affect the same molecular pathway (66) indicating that these miRNAs could be involved and potentially serve as common biomarkers for diagnosing different models of liver disease associated with AATD. Therefore, the present study suggests that liver PASD and fibrosis in AATD individuals share plasma EV-associated miRNA profiles. In addition, we observed that most circulating DEMs were correlated with pathological changes in the liver of AATD individuals.

Liver biopsy is the currently used standard diagnostic method for various liver diseases. However, the difficulty, invasiveness, and risk related to sampling have promoted the exploration of the plasma derived EV as a potential alternative in these pathologic scenarios. Our studies evaluated the effectiveness of plasma EV-derived miRNAs in identifying AATD individuals with liver disease (PASD 2-3) as compared to those without liver disease (PASD 0-1). Our qPCR analysis confirmed the levels of plasma EV-associated miR-23a-3p were significantly elevated in AATD individuals. Furthermore, we observed that miR-23a-3p levels were significantly higher in AATD individuals with liver disease compared to those without liver disease. High-throughput sequencing analysis has previously identified miR-23a-3p as one of the most abundant EV-associated miRNAs released by hepatocytes with endoplasmic reticulum stress, which is negatively correlated with hepatocyte viability (67). Additionally, overexpression of miR-23a has been shown to damage mitochondria in vitro (68) and promote triglyceride accumulation in hepatocytes (69). Our data indicating overexpression of miR-23a in AATD plasma EV is the first report supporting our previous findings of critical role for hepatic ER stress (70), lipid accumulation (71), and mitochondrial dysfunction (72, 73) in the progression of AATD-mediated liver disease. This result provides further evidence that analyzing plasma EV-associated levels of miR-23a-3p could provide insights into the mechanisms linked to AATD-mediated liver disease.

Numerous studies have shown that the combination of multiple miRNAs can obtain higher discrimination than a single miRNA biomarker (74). Therefore, we selected 16 miRNAs which exhibited the AUC of 0.7 or higher which were differentially expressed in these two groups of AATD individuals. Then, we tightened the range of candidate miRNAs into 6 miRNAs biologically associated with liver disease to maximize the success rate of multiple EV biomarker verification. We found that three-biomarker panels including combinations of miR-223-3p, miR-23a-3p, miR-15b-5p, let-7a-5p and let-7f-5p could identify liver disease within AATD individuals with a sensitivity of 100% and a specificity of 83.3% (AUC> 0.93) in our training batch. These three-biomarker panels represent a sensitivity of 66-77% and a specificity of 60% (AUC> 0.73) in the validation batch (Fig. 6C). We also observed that a four-biomarker panel including miR-223-3p, miR-374a-3p, miR-23a-3p, and miR-15b-5p presented sensitivity and specificity of 100% (AUC=1) for identifying liver disease within AATD individuals in our training batch and an AUC of 0.75 in the validation batch (Fig. 6D). These panels represent an innovative way to monitor liver disease status in the AATD population. ER stress, mitochondrial dysfunction, and lipotoxicity are main hallmarks of AATD mediated liver disease (75). In this regard, miR- 223, miR-23a, miRNA-15b, and miR374a have been previously shown to be correlated with liver fibrosis, regulating lipid and cholesterol homeostasis (69, 76–78). Furthermore, miRNA-15b has been reported to regulate ATP levels and mitochondrial ROS production (79, 80), while miR223-3p increases mitophagy in cells with dysfunctional mitochondria (81). Interestingly, serum levels of miR-15b, miR-23a, miR-374a, and let-7a have been recently reported to have great potential as diagnostic biomarkers of nonalcoholic liver fibrosis (82). Altogether, these results provide the basis for further research of how the pathways related to these miRNAs are involved in the pathogenesis of AATD liver disease and their impact on disease progression. An unexpected finding of this study is that miR- 425-5p had diagnostic power, with an AUC of 0.79, sensitivity of 66% and specificity of 94% for detecting AATD individuals with liver disease in the validation set. Our results also indicated that miR-425-5p has AUC of 0.62, sensitivity of 90%, and specificity of 50% for detecting AATD individuals with liver disease in the training set. miR-425-5p has been previously reported to regulate liver inflammation in inflammatory liver diseases (83, 84). Recent evidence also shows that over expression of miR-425-5p is significantly associated with poor prognosis of hepatocellular carcinoma patients whereas knockdown of miR-425-5p inhibits cell proliferation and migration (85). Our data clearly indicated that miR-425-5p expression was significantly decreased in AATD individuals. Given that AATD individuals may have an increased risk of developing hepatocellular carcinoma (86) decreased levels of plasma EV-associated miR-425-5p may suggest a defense mechanism to prevent cancer in this patient’s population.

Herein, we also investigated the mRNA transcriptome of AATD liver tissues at different stages of liver disease and observed that the liver transcriptome and associated functional annotations were largely overlapping in AATD individuals with different stages of liver disease. Our AATD liver RNA-seq data revealed that during the progression of liver disease following hepatic accumulation of PASD globules, 273 genes show significant upregulation, and 12 genes were downregulated. These genes were involved in various cellular and molecular mechanisms associated with development of fibrosis, mostly related to extracellular matrix remodeling. A diverse array of mechanisms and causal genes are implicated in the pathogenesis of liver fibrosis. The fibrotic phenotypes include the molecular changes in hepatic stellate cells (HSC) in response to persistent inflammatory signals which have undergone ‘‘activation’’ to pro-fibrogenic myofibroblasts (87). Interestingly, activated HSC express several neuronal and muscle cell markers such as MyoD, α–smooth muscle actin (α-SMA), and MEF2 (88). The KEGG enrichment analysis revealed that the key targets of DEGs during progression of AATD mediated liver disease were mainly enriched in the cardiomyopathy, motor proteins, oxytocin signaling pathway, and glucagon signaling pathway. Oxytocin, with its anti-inflammatory and antioxidant effects on the liver (89), has been demonstrated to play role in hepatic regeneration (90). Changes in the glucagon signaling pathway have also been shown to be associated with fibrotic progression of the liver (91). The G.O. enrichment analysis revealed that AATD livers undergo transcriptomic changes mostly involved in the regulation of ECM remodeling, regulation of muscle cell proliferation, regulation of smooth muscle cell proliferation and hepatic metabolism. IPA also predicted that during the progression of AATD-mediated liver disease, many dysregulated genes in the AATD liver are involved in the mechanisms related to hepatic stellate cell activation. The extensive regulation of extracellular matrix-associated genes (92) also indicates that major transcriptome changes in AATD-mediated liver disease are attributed to the histopathological evidence of fibrosis in these individuals. Therefore, the overlap between liver transcriptome profiles associated with the AATD-mediated liver disease and fibrosis support the concept that a sequence of interacting molecular mechanisms and signaling pathways drives the transition from hepatic PASD accumulation to liver fibrosis.

It has been evidenced that miRNAs are epigenetic gene expression modulators, inhibiting the expression of target genes. Although there are a lot of miRNAs playing roles in the progression of liver fibrosis, there is limited evidence on miRNA-mRNA interactions and their regulatory mechanisms involved in liver fibrogenesis (93). Therefore, in the present study we have combined RNA-sequencing data from AATD liver tissues, with plasma EV-miRNA expression data to systematically identify the genes which may have been regulated by plasma EV-miRNAs in AATD individuals. By sequencing miRNAs and mRNAs from the same individuals, we have a unique possibility to investigate how miRNA and mRNAs interact to regulate each other (94). Based on our analysis, liver mRNA and plasma EV-miRNA expression data in AATD individuals share many common features, especially those related to mitochondrial dysfunction, liver cirrhosis, and liver fibrogenesis. We observed that miR-3064-3p, let-7a-5p, and miR-16-5p target liver mRNAs which have dysregulated expression during progression of AATD liver disease, including MAPK8, PRKAG1, and PRKAR2A. As a canonical pathway involved in lipid metabolism, PPAR activation expends anti-inflammatory and anti-fibrosis effects due to its extended protection for hepatocytes, liver sinusoidal endothelial cells, and HSCs (95). Furthermore, multiple studies also provide evidence that in the liver, Mitogen-Activated Protein Kinases (MAPKs) play a proinflammatory role by increasing the secretion of inflammatory cytokines leading to the activation and infiltration of immune cells into the liver tissue (96). MAPKs, including protein kinase A family, have been shown to be involved in activation of hepatic stellate cells (96, 97) liver steatosis, and fibrosis (98, 99). Therefore, our study provides new insight that alteration of the MAPK signaling pathway mediated by miRNAs contributes to liver disease in AATD individuals.

## CONCLUSION

In conclusion, this study suggested that AATD individuals retained a specific plasma EV derived miRNA profile compared with healthy controls. These miRNAs mostly play roles in influencing molecular and cellular processes in tissue remodeling, fibrosis, and inflammation. While there is no biomarker study for AATD mediated liver disease, here we suggest that AATD individuals with liver disease have a distinct plasma EV-associated miRNA profile as compared to individuals without liver disease. In this study, we illustrated the unique properties of six plasma EV-associated miRNAs (let-7a-5p, let-7f-5p, miR-223- 3p, miR-23a-3p, miR-374-3p, and miR-15b-5p) and their combinations, which may be a new promising sensitive biomarker category for PASD, fibrosis progression, and for detection of early and small changes in AATD-mediated liver disease. They can serve to identify patients at risk to develop advanced liver disease, while being a great tool for studying the effects of anti-fibrotic drugs by shortening the duration of clinical trials due to easy and noninvasive detection of small changes (100). Furthermore, we identified the mRNA expression profiles of liver tissues during progression of AATD-mediated liver disease. DEGs during progression of AATD mediated liver disease are mostly involve in metabolic changes, ECM remodeling, and liver fibrogenesis consistent with our clinical observations. Our study for the first time also described miRNA and mRNA expression profiles associated with AATD-mediated liver disease and identified novel miRNA-mRNA interaction networks and signaling pathways. All these might be helpful to understand pathophysiologic changes of the AATD liver or AATD-related fibrogenesis at the transcriptional and post-transcriptional level.

## LIMITATIONS

Although our results suggest the possibility of plasma EV miRNAs to be used as part of a novel biomarker panel in AATD individuals with liver disease, we acknowledge that our study has several limitations, such as the number of individuals included. However, even with this limited number of subjects, significant changes could be found in the miRNA levels of plasma circulating miRNAs. Analysis of larger cohorts of AATD patients will increase power and can perhaps provide enough evidence to include one or more of the proposed miRNAs in a future diagnostic panel for early stage AATD-mediated liver disease. Furthermore, longitudinal human data would eventually be necessary to further support our findings. The potential candidate targets for the diagnosis and therapeutic approach of AATD-mediated liver disease provided in this study also need further elucidation.

## AUTHOR CONTRIBUTION

R.O.: acquisition of data, technical support, review & editing. Z. H.: Data curation; Formal analysis, review & editing. B. P.: acquisition of data, Writing – review & editing. V. C.: acquisition of data. H. Z.: Formal analysis, review & editing. J.W.: acquisition of data. M.W.: Data curation; Formal analysis. M.A.: Data curation, Formal analysis. M.H.: Writing – review & editing. M.B.: Conceptualization, Supervision. N.K.: Conceptualization, Funding acquisition, Formal analysis, Data curation, Supervision, Writing the original draft, review & editing.

## GEOLOCATION INFORMATION

29.64079685199175, -82.34289418000674

## ACKNOWLEDGMENT

We would like to thank Ashish Sharma, Shannon Holiday, and Asha Rani for their advice and suggestions on this work and members of our laboratories for their contribution to various aspects of our research. RNA sequencing, Small RNA sequencing and data analysis were performed at the Interdisciplinary Center for Biotechnology Research in the University of Florida’s premium core research facility under the supervision of Drs. Yanping Zhang and Tongjun Gu.

## DISCLOUSURE STATMENT

No potential conflict of interest was reported by the authors of this manuscript.

## DATA SHARING STATEMENT

The data that supports the findings of this study are available in the supplementary material of this article

---

**Figure S1.**
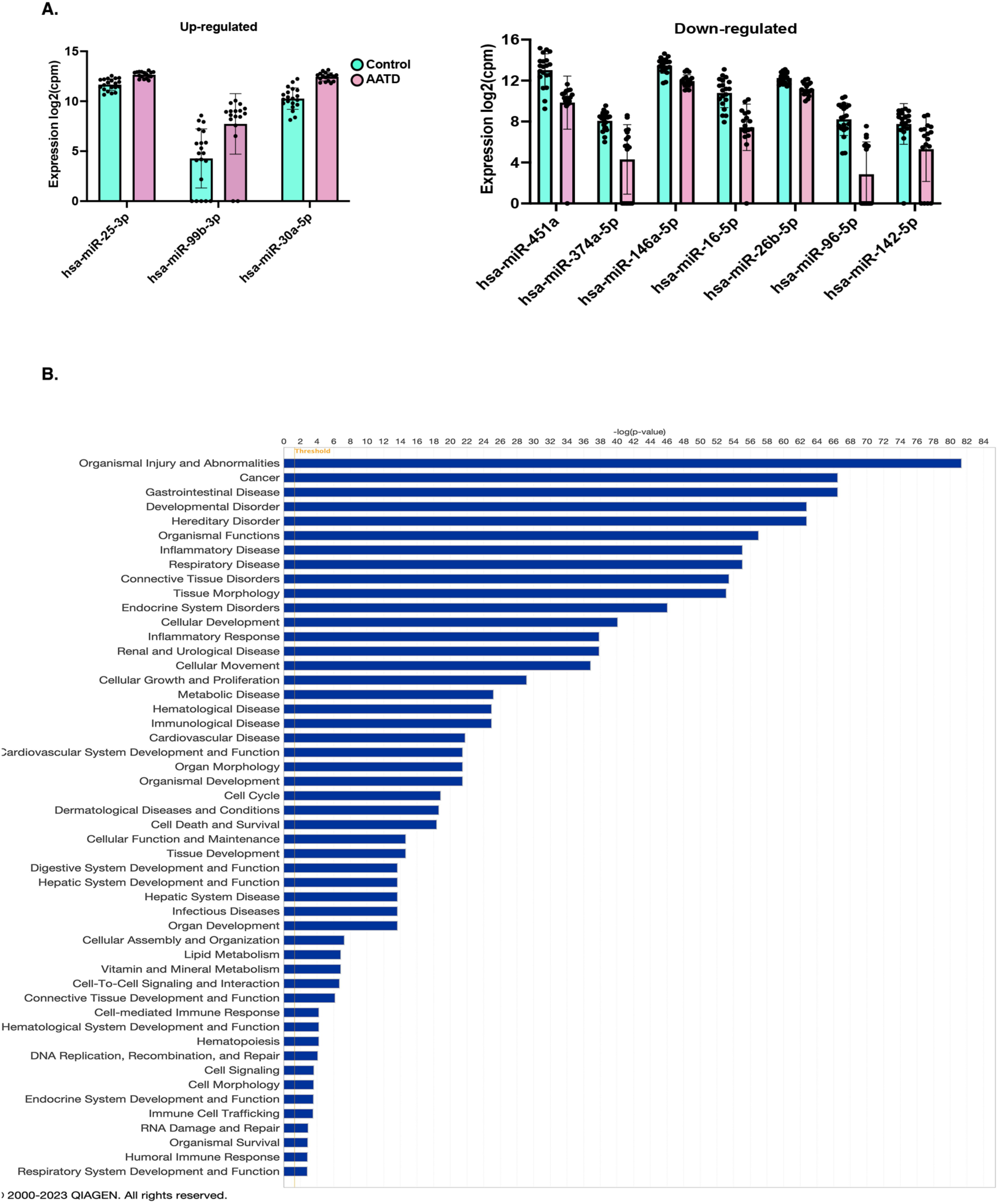
Top 10 differentially expressed miRNAs in plasma Extracellular Vesicles (EV) from alpha-1 antitrypsin deficient (AATD) individuals. **A.** Graph bars indicating upregulated and downregulated differentially expressed miRNAs from our plasma EV fractions miRNA dataset. **B.** A bar plot shows the numbers of diseases and abnormalities associated with top 10 differentially expressed miRNAs (DEMs) in our plasma EV fractions miRNA dataset.

**Figure S2.**
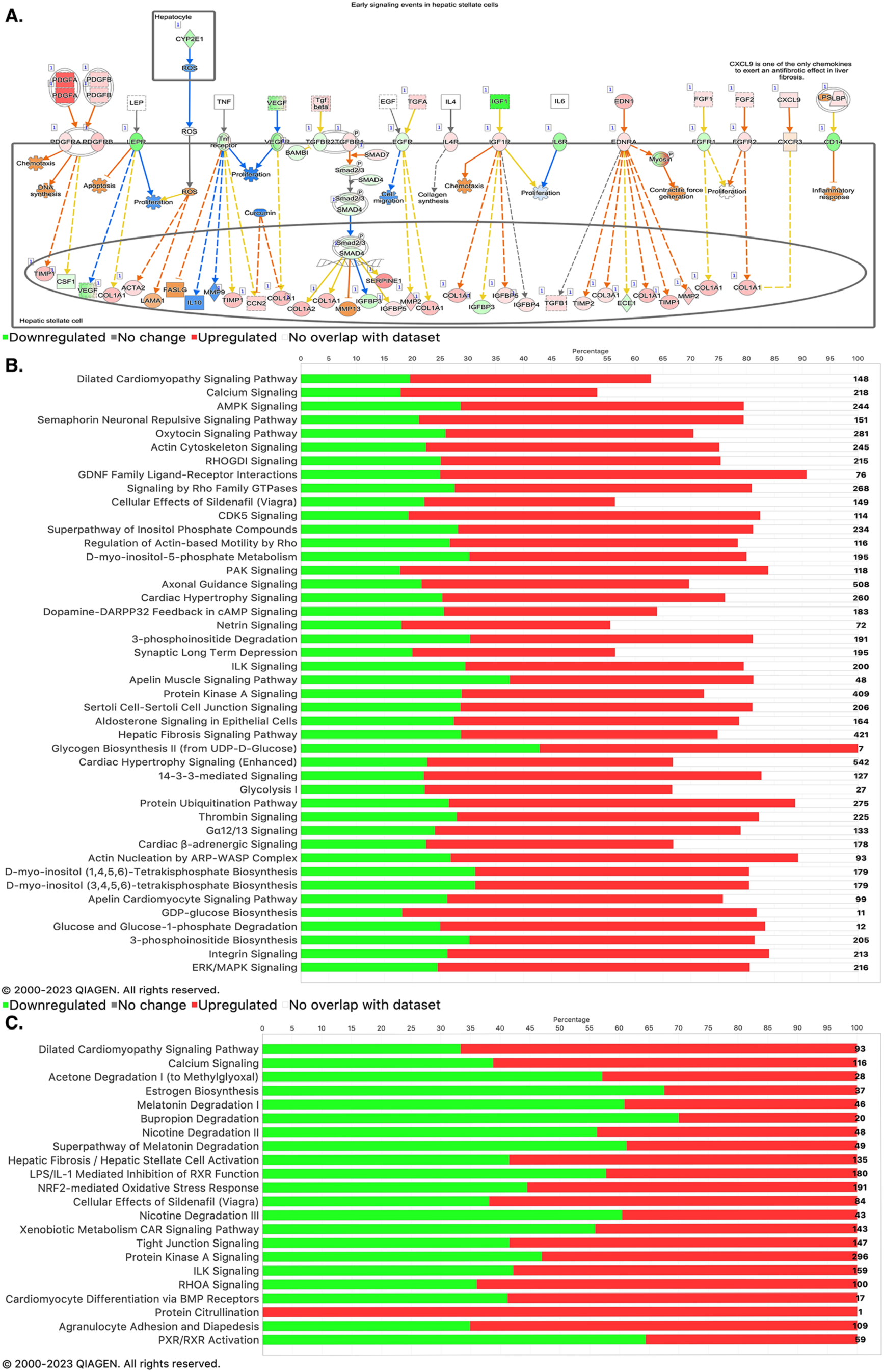
Pathway identification and functional enrichment of liver transcriptomics in alpha-1 antitrypsin deficient (AATD) individuals. **A.** Ingenuity Pathway Analysis (IPA) shows signaling events in hepatic stellate cells during progression of AATD-mediated liver disease. Red indicates upregulated miRNAs and green indicates downregulated miRNAs from our dataset. Orange color indicates upregulation and blue indicates downregulation of target genes, predicted by IPA software. **B.** Canonical pathways related to differentially expressed genes in the liver of AATD individuals with PASD 2 as compared to those livers with PASD 0. **C.** Canonical pathways related to the 55 differentially expressed genes when comparing livers with PASD 3 to those with PASD 0 as predicted by IPA software.

## REFERENCES

1. Janciauskiene SM, Bals R, Koczulla R, Vogelmeier C, Kohnlein T, Welte T. The discovery of alpha1-antitrypsin and its role in health and disease. Respir Med 2011; 105: 1129–1139.

2. Lindblad D, Blomenkamp K, Teckman J. Alpha-1-antitrypsin mutant Z protein content in individual hepatocytes correlates with cell death in a mouse model. Hepatology 2007; 46: 1228–1235.

3. Crowther DC, Belorgey D, Miranda E, Kinghorn KJ, Sharp LK, Lomas DA. Practical genetics: alpha-1-antitrypsin deficiency and the serpinopathies. Eur J Hum Genet 2004; 12: 167–172.

4. Khodayari N, Oshins R, Holliday LS, Clark V, Xiao Q, Marek G, Mehrad B, Brantly M. Alpha-1 antitrypsin deficient individuals have circulating extracellular vesicles with profibrogenic cargo. Cell Commun Signal 2020; 18: 140.

5. Perlmutter DH. Current and Emerging Treatments for Alpha-1 Antitrypsin Deficiency. Gastroenterol Hepatol (N Y) 2016; 12: 446–448.

6. Zamora MR, Ataya A. Lung and liver transplantation in patients with alpha-1 antitrypsin deficiency. Ther Adv Chronic Dis 2021; 12_suppl: 20406223211002988.

7. El-Serag HB. Hepatocellular carcinoma and hepatitis C in the United States. Hepatology 2002; 36: S74–83.

8. Sun M, Kisseleva T. Reversibility of liver fibrosis. Clin Res Hepatol Gastroenterol 2015; 39 Suppl 1: S60–63.

9. Clark VC, Marek G, Liu C, Collinsworth A, Shuster J, Kurtz T, Nolte J, Brantly M. Clinical and histologic features of adults with alpha-1 antitrypsin deficiency in a non-cirrhotic cohort. J Hepatol 2018; 69: 1357–1364.

10. Chang Y, Han JA, Kang SM, Jeong SW, Ryu T, Park HS, Yoo JJ, Lee SH, Kim SG, Kim YS, Kim HS, Jin SY, Ryu S, Jang JY. Clinical impact of serum exosomal microRNA in liver fibrosis. PLoS One 2021; 16: e0255672.

11. Abdi W, Millan JC, Mezey E. Sampling variability on percutaneous liver biopsy. Arch Intern Med 1979; 139: 667–669.

12. Abels ER, Breakefield XO. Introduction to Extracellular Vesicles: Biogenesis, RNA Cargo Selection, Content, Release, and Uptake. Cell Mol Neurobiol 2016; 36: 301–312.

13. Sung S, Kim J, Jung Y. Liver-Derived Exosomes and Their Implications in Liver Pathobiology. Int J Mol Sci 2018; 19.

14. Jiao Y, Xu P, Shi H, Chen D, Shi H. Advances on liver cell-derived exosomes in liver diseases. J Cell Mol Med 2021; 25: 15–26.

15. Yanez-Mo M, Siljander PR, Andreu Z, Zavec AB, Borras FE, Buzas EI, Buzas K, Casal E, Cappello F, Carvalho J, Colas E, Cordeiro-da Silva A, Fais S, Falcon-Perez JM, Ghobrial IM, Giebel B, Gimona M, Graner M, Gursel I, Gursel M, Heegaard NH, Hendrix A, Kierulf P, Kokubun K, Kosanovic M, Kralj-Iglic V, Kramer-Albers EM, Laitinen S, Lasser C, Lener T, Ligeti E, Line A, Lipps G, Llorente A, Lotvall J, Mancek-Keber M, Marcilla A, Mittelbrunn M, Nazarenko I, Nolte-’t Hoen EN, Nyman TA, O’Driscoll L, Olivan M, Oliveira C, Pallinger E, Del Portillo HA, Reventos J, Rigau M, Rohde E, Sammar M, Sanchez-Madrid F, Santarem N, Schallmoser K, Ostenfeld MS, Stoorvogel W, Stukelj R, Van der Grein SG, Vasconcelos MH, Wauben MH, De Wever O. Biological properties of extracellular vesicles and their physiological functions. J Extracell Vesicles 2015; 4: 27066.

16. Jiao Y, Lu W, Xu P, Shi H, Chen D, Chen Y, Shi H, Ma Y. Hepatocyte-derived exosome may be as a biomarker of liver regeneration and prognostic valuation in patients with acute-on-chronic liver failure. Hepatol Int 2021; 15: 957–969.

17. Roderburg C, Luedde T. Circulating microRNAs as markers of liver inflammation, fibrosis and cancer. J Hepatol 2014; 61: 1434–1437.

18. Bala S, Petrasek J, Mundkur S, Catalano D, Levin I, Ward J, Alao H, Kodys K, Szabo G. Circulating microRNAs in exosomes indicate hepatocyte injury and inflammation in alcoholic, drug-induced, and inflammatory liver diseases. Hepatology 2012; 56: 1946–1957.

19. Masyuk AI, Masyuk TV, Larusso NF. Exosomes in the pathogenesis, diagnostics and therapeutics of liver diseases. J Hepatol 2013; 59: 621–625.

20. Cortez MA, Bueso-Ramos C, Ferdin J, Lopez-Berestein G, Sood AK, Calin GA. MicroRNAs in body fluids--the mix of hormones and biomarkers. Nat Rev Clin Oncol 2011; 8: 467–477.

21. Bandopadhyay M, Bharadwaj M. Exosomal miRNAs in hepatitis B virus related liver disease: a new hope for biomarker. Gut Pathog 2020; 12: 23.

22. Cieri UR. Determination of ergotamine tartrate in tablets by liquid chromatography with fluorescence detection. J Assoc Off Anal Chem 1987; 70: 538–540.

23. Lynoe N, Sandlund M, Dahlqvist G, Jacobsson L. Informed consent: study of quality of information given to participants in a clinical trial. BMJ 1991; 303: 610–613.

24. Brantly ML, Wittes JT, Vogelmeier CF, Hubbard RC, Fells GA, Crystal RG. Use of a highly purified alpha 1-antitrypsin standard to establish ranges for the common normal and deficient alpha 1-antitrypsin phenotypes. Chest 1991; 100: 703–708.

25. Thery C, Witwer KW, Aikawa E, Alcaraz MJ, Anderson JD, Andriantsitohaina R, Antoniou A, Arab T, Archer F, Atkin-Smith GK, Ayre DC, Bach JM, Bachurski D, Baharvand H, Balaj L, Baldacchino S, Bauer NN, Baxter AA, Bebawy M, Beckham C, Bedina Zavec A, Benmoussa A, Berardi AC, Bergese P, Bielska E, Blenkiron C, Bobis-Wozowicz S, Boilard E, Boireau W, Bongiovanni A, Borras FE, Bosch S, Boulanger CM, Breakefield X, Breglio AM, Brennan MA, Brigstock DR, Brisson A, Broekman ML, Bromberg JF, Bryl-Gorecka P, Buch S, Buck AH, Burger D, Busatto S, Buschmann D, Bussolati B, Buzas EI, Byrd JB, Camussi G, Carter DR, Caruso S, Chamley LW, Chang YT, Chen C, Chen S, Cheng L, Chin AR, Clayton A, Clerici SP, Cocks A, Cocucci E, Coffey RJ, Cordeiro-da-Silva A, Couch Y, Coumans FA, Coyle B, Crescitelli R, Criado MF, D’Souza-Schorey C, Das S, Datta Chaudhuri A, de Candia P, De Santana EF, De Wever O, Del Portillo HA, Demaret T, Deville S, Devitt A, Dhondt B, Di Vizio D, Dieterich LC, Dolo V, Dominguez Rubio AP, Dominici M, Dourado MR, Driedonks TA, Duarte FV, Duncan HM, Eichenberger RM, Ekstrom K, El Andaloussi S, Elie-Caille C, Erdbrugger U, Falcon-Perez JM, Fatima F, Fish JE, Flores-Bellver M, Forsonits A, Frelet-Barrand A, Fricke F, Fuhrmann G, Gabrielsson S, Gamez-Valero A, Gardiner C, Gartner K, Gaudin R, Gho YS, Giebel B, Gilbert C, Gimona M, Giusti I, Goberdhan DC, Gorgens A, Gorski SM, Greening DW, Gross JC, Gualerzi A, Gupta GN, Gustafson D, Handberg A, Haraszti RA, Harrison P, Hegyesi H, Hendrix A, Hill AF, Hochberg FH, Hoffmann KF, Holder B, Holthofer H, Hosseinkhani B, Hu G, Huang Y, Huber V, Hunt S, Ibrahim AG, Ikezu T, Inal JM, Isin M, Ivanova A, Jackson HK, Jacobsen S, Jay SM, Jayachandran M, Jenster G, Jiang L, Johnson SM, Jones JC, Jong A, Jovanovic-Talisman T, Jung S, Kalluri R, Kano SI, Kaur S, Kawamura Y, Keller ET, Khamari D, Khomyakova E, Khvorova A, Kierulf P, Kim KP, Kislinger T, Klingeborn M, Klinke DJ, 2nd, Kornek M, Kosanovic MM, Kovacs AF, Kramer-Albers EM, Krasemann S, Krause M, Kurochkin IV, Kusuma GD, Kuypers S, Laitinen S, Langevin SM, Languino LR, Lannigan J, Lasser C, Laurent LC, Lavieu G, Lazaro-Ibanez E, Le Lay S, Lee MS, Lee YXF, Lemos DS, Lenassi M, Leszczynska A, Li IT, Liao K, Libregts SF, Ligeti E, Lim R, Lim SK, Line A, Linnemannstons K, Llorente A, Lombard CA, Lorenowicz MJ, Lorincz AM, Lotvall J, Lovett J, Lowry MC, Loyer X, Lu Q, Lukomska B, Lunavat TR, Maas SL, Malhi H, Marcilla A, Mariani J, Mariscal J, Martens-Uzunova ES, Martin-Jaular L, Martinez MC, Martins VR, Mathieu M, Mathivanan S, Maugeri M, McGinnis LK, McVey MJ, Meckes DG, Jr., Meehan KL, Mertens I, Minciacchi VR, Moller A, Moller Jorgensen M, Morales-Kastresana A, Morhayim J, Mullier F, Muraca M, Musante L, Mussack V, Muth DC, Myburgh KH, Najrana T, Nawaz M, Nazarenko I, Nejsum P, Neri C, Neri T, Nieuwland R, Nimrichter L, Nolan JP, Nolte-’t Hoen EN, Noren Hooten N, O’Driscoll L, O’Grady T, O’Loghlen A, Ochiya T, Olivier M, Ortiz A, Ortiz LA, Osteikoetxea X, Ostergaard O, Ostrowski M, Park J, Pegtel DM, Peinado H, Perut F, Pfaffl MW, Phinney DG, Pieters BC, Pink RC, Pisetsky DS, Pogge von Strandmann E, Polakovicova I, Poon IK, Powell BH, Prada I, Pulliam L, Quesenberry P, Radeghieri A, Raffai RL, Raimondo S, Rak J, Ramirez MI, Raposo G, Rayyan MS, Regev-Rudzki N, Ricklefs FL, Robbins PD, Roberts DD, Rodrigues SC, Rohde E, Rome S, Rouschop KM, Rughetti A, Russell AE, Saa P, Sahoo S, Salas-Huenuleo E, Sanchez C, Saugstad JA, Saul MJ, Schiffelers RM, Schneider R, Schoyen TH, Scott A, Shahaj E, Sharma S, Shatnyeva O, Shekari F, Shelke GV, Shetty AK, Shiba K, Siljander PR, Silva AM, Skowronek A, Snyder OL, 2nd, Soares RP, Sodar BW, Soekmadji C, Sotillo J, Stahl PD, Stoorvogel W, Stott SL, Strasser EF, Swift S, Tahara H, Tewari M, Timms K, Tiwari S, Tixeira R, Tkach M, Toh WS, Tomasini R, Torrecilhas AC, Tosar JP, Toxavidis V, Urbanelli L, Vader P, van Balkom BW, van der Grein SG, Van Deun J, van Herwijnen MJ, Van Keuren-Jensen K, van Niel G, van Royen ME, van Wijnen AJ, Vasconcelos MH, Vechetti IJ, Jr., Veit TD, Vella LJ, Velot E, Verweij FJ, Vestad B, Vinas JL, Visnovitz T, Vukman KV, Wahlgren J, Watson DC, Wauben MH, Weaver A, Webber JP, Weber V, Wehman AM, Weiss DJ, Welsh JA, Wendt S, Wheelock AM, Wiener Z, Witte L, Wolfram J, Xagorari A, Xander P, Xu J, Yan X, Yanez-Mo M, Yin H, Yuana Y, Zappulli V, Zarubova J, Zekas V, Zhang JY, Zhao Z, Zheng L, Zheutlin AR, Zickler AM, Zimmermann P, Zivkovic AM, Zocco D, Zuba-Surma EK. Minimal information for studies of extracellular vesicles 2018 (MISEV2018): a position statement of the International Society for Extracellular Vesicles and update of the MISEV2014 guidelines. J Extracell Vesicles 2018; 7: 1535750.

26. Kechin A, Boyarskikh U, Kel A, Filipenko M. cutPrimers: A New Tool for Accurate Cutting of Primers from Reads of Targeted Next Generation Sequencing. J Comput Biol 2017; 24: 1138–1143.

27. Langdon WB. Performance of genetic programming optimised Bowtie2 on genome comparison and analytic testing (GCAT) benchmarks. BioData Min 2015; 8: 1.

28. Langmead B, Salzberg SL. Fast gapped-read alignment with Bowtie 2. Nat Methods 2012; 9: 357–359.

29. Anders S, Pyl PT, Huber W. HTSeq--a Python framework to work with high-throughput sequencing data. Bioinformatics 2015; 31: 166–169.

30. Kozomara A, Griffiths-Jones S. miRBase: annotating high confidence microRNAs using deep sequencing data. Nucleic Acids Res 2014; 42: D68–73.

31. Bolger AM, Lohse M, Usadel B. Trimmomatic: a flexible trimmer for Illumina sequence data. Bioinformatics 2014; 30: 2114–2120.

32. Phan L, Hsu J, Tri LQ, Willi M, Mansour T, Kai Y, Garner J, Lopez J, Busby B. dbVar structural variant cluster set for data analysis and variant comparison. F1000Res 2016; 5: 673.

33. Li B, Dewey CN. RSEM: accurate transcript quantification from RNA-Seq data with or without a reference genome. BMC Bioinformatics 2011; 12: 323.

34. Khodayari N, Oshins R, Aranyos AM, Duarte S, Mostofizadeh S, Lu Y, Brantly M. Characterization of hepatic inflammatory changes in a C57BL/6J mouse model of alpha1-antitrypsin deficiency. Am J Physiol Gastrointest Liver Physiol 2022; 323: G594–G608.

35. Evangelista JE, Xie Z, Marino GB, Nguyen N, Clarke DJB, Ma’ayan A. Enrichr-KG: bridging enrichment analysis across multiple libraries. Nucleic Acids Res 2023.

36. Wu T, Hu E, Xu S, Chen M, Guo P, Dai Z, Feng T, Zhou L, Tang W, Zhan L, Fu X, Liu S, Bo X, Yu G. clusterProfiler 4.0: A universal enrichment tool for interpreting omics data. Innovation (Camb) 2021; 2: 100141.

37. Ye J, Fang L, Zheng H, Zhang Y, Chen J, Zhang Z, Wang J, Li S, Li R, Bolund L, Wang J. WEGO: a web tool for plotting GO annotations. Nucleic Acids Res 2006; 34: W293–297.

38. Chen XM. MicroRNA signatures in liver diseases. World J Gastroenterol 2009; 15: 1665–1672.

39. Robinson MD, Oshlack A. A scaling normalization method for differential expression analysis of RNA-seq data. Genome Biol 2010; 11: R25.

40. Robinson MD, McCarthy DJ, Smyth GK. edgeR: a Bioconductor package for differential expression analysis of digital gene expression data. Bioinformatics 2010; 26: 139–140.

41. Matsuura K, Aizawa N, Enomoto H, Nishiguchi S, Toyoda H, Kumada T, Iio E, Ito K, Ogawa S, Isogawa M, Alter HJ, Tanaka Y. Circulating let-7 Levels in Serum Correlate With the Severity of Hepatic Fibrosis in Chronic Hepatitis C. Open Forum Infect Dis 2018; 5: ofy268.

42. Wen Y, Han J, Chen J, Dong J, Xia Y, Liu J, Jiang Y, Dai J, Lu J, Jin G, Han J, Wei Q, Shen H, Sun B, Hu Z. Plasma miRNAs as early biomarkers for detecting hepatocellular carcinoma. Int J Cancer 2015; 137: 1679–1690.

43. Gharib AF, Eed EM, Khalifa AS, Raafat N, Shehab-Eldeen S, Alwakeel HR, Darwiesh E, Essa A. Value of Serum miRNA-96-5p and miRNA-99a-5p as Diagnostic Biomarkers for Hepatocellular Carcinoma. Int J Gen Med 2022; 15: 2427–2436.

44. Song LY, Ma YT, Wu CF, Wang CJ, Fang WJ, Liu SK. MicroRNA-195 Activates Hepatic Stellate Cells In Vitro by Targeting Smad7. Biomed Res Int 2017; 2017: 1945631.

45. Shao M, Xu Q, Wu Z, Chen Y, Shu Y, Cao X, Chen M, Zhang B, Zhou Y, Yao R, Shi Y, Bu H. Exosomes derived from human umbilical cord mesenchymal stem cells ameliorate IL-6-induced acute liver injury through miR-455-3p. Stem Cell Res Ther 2020; 11: 37.

46. Guo Y, Xiong Y, Sheng Q, Zhao S, Wattacheril J, Flynn CR. A micro-RNA expression signature for human NAFLD progression. J Gastroenterol 2016; 51: 1022–1030.

47. Liu YM, Ma JH, Zeng QL, Lv J, Xie XH, Pan YJ, Yu ZJ. MiR-19a Affects Hepatocyte Autophagy via Regulating lncRNA NBR2 and AMPK/PPARalpha in D-GalN/Lipopolysaccharide-Stimulated Hepatocytes. J Cell Biochem 2018; 119: 358–365.

48. Lakner AM, Steuerwald NM, Walling TL, Ghosh S, Li T, McKillop IH, Russo MW, Bonkovsky HL, Schrum LW. Inhibitory effects of microRNA 19b in hepatic stellate cell-mediated fibrogenesis. Hepatology 2012; 56: 300–310.

49. Yao X, Zhang H, Liu Y, Liu X, Wang X, Sun X, Cheng Y. miR-99b-3p promotes hepatocellular carcinoma metastasis and proliferation by targeting protocadherin 19. Gene 2019; 698: 141–149.

50. Li WF, Dai H, Ou Q, Zuo GQ, Liu CA. Overexpression of microRNA-30a-5p inhibits liver cancer cell proliferation and induces apoptosis by targeting MTDH/PTEN/AKT pathway. Tumour Biol 2016; 37: 5885–5895.

51. Cheng B, Ding F, Huang CY, Xiao H, Fei FY, Li J. Role of miR-16-5p in the proliferation and metastasis of hepatocellular carcinoma. Eur Rev Med Pharmacol Sci 2019; 23: 137–145.

52. Messner CJ, Schmidt S, Ozkul D, Gaiser C, Terracciano L, Krahenbuhl S, Suter-Dick L. Identification of miR-199a-5p, miR-214-3p and miR-99b-5p as Fibrosis-Specific Extracellular Biomarkers and Promoters of HSC Activation. Int J Mol Sci 2021; 22.

53. Jiang XP, Ai WB, Wan LY, Zhang YQ, Wu JF. The roles of microRNA families in hepatic fibrosis. Cell Biosci 2017; 7: 34.

54. Wang C, Wang X, Su Z, Fei H, Liu X, Pan Q. MiR-25 promotes hepatocellular carcinoma cell growth, migration and invasion by inhibiting RhoGDI1. Oncotarget 2015; 6: 36231–36244.

55. Damanti CC, Gaffo E, Lovisa F, Garbin A, Di Battista P, Gallingani I, Tosato A, Pillon M, Carraro E, Mascarin M, Elia C, Biffi A, Bortoluzzi S, Mussolin L. MiR-26a-5p as a Reference to Normalize MicroRNA qRT-PCR Levels in Plasma Exosomes of Pediatric Hematological Malignancies. Cells 2021; 10.

56. Zhang Y, Han T, Feng D, Li J, Wu M, Peng X, Wang B, Zhan X, Fu P. Screening of non-invasive miRNA biomarker candidates for metastasis of gastric cancer by small RNA sequencing of plasma exosomes. Carcinogenesis 2020; 41: 582–590.

57. Mateescu B, Kowal EJ, van Balkom BW, Bartel S, Bhattacharyya SN, Buzas EI, Buck AH, de Candia P, Chow FW, Das S, Driedonks TA, Fernandez-Messina L, Haderk F, Hill AF, Jones JC, Van Keuren-Jensen KR, Lai CP, Lasser C, Liegro ID, Lunavat TR, Lorenowicz MJ, Maas SL, Mager I, Mittelbrunn M, Momma S, Mukherjee K, Nawaz M, Pegtel DM, Pfaffl MW, Schiffelers RM, Tahara H, Thery C, Tosar JP, Wauben MH, Witwer KW, Nolte-’t Hoen EN. Obstacles and opportunities in the functional analysis of extracellular vesicle RNA - an ISEV position paper. J Extracell Vesicles 2017; 6: 1286095.

58. Min L, Zhu S, Chen L, Liu X, Wei R, Zhao L, Yang Y, Zhang Z, Kong G, Li P, Zhang S. Evaluation of circulating small extracellular vesicles derived miRNAs as biomarkers of early colon cancer: a comparison with plasma total miRNAs. J Extracell Vesicles 2019; 8: 1643670.

59. Rudnick DA, Perlmutter DH. Alpha-1-antitrypsin deficiency: a new paradigm for hepatocellular carcinoma in genetic liver disease. Hepatology 2005; 42: 514–521.

60. Li SP, Lin ZX, Jiang XY, Yu XY. Exosomal cargo-loading and synthetic exosome-mimics as potential therapeutic tools. Acta Pharmacol Sin 2018; 39: 542–551.

61. Ge S, Wang X, Xie J, Yi X, Liu F. Deep sequencing analysis of microRNA expression in porcine serum-induced hepatic fibrosis rats. Ann Hepatol 2014; 13: 439–449.

62. Khalil AM, Guttman M, Huarte M, Garber M, Raj A, Rivea Morales D, Thomas K, Presser A, Bernstein BE, van Oudenaarden A, Regev A, Lander ES, Rinn JL. Many human large intergenic noncoding RNAs associate with chromatin-modifying complexes and affect gene expression. Proc Natl Acad Sci U S A 2009; 106: 11667–11672.

63. Bian EB, Wang YY, Yang Y, Wu BM, Xu T, Meng XM, Huang C, Zhang L, Lv XW, Xiong ZG, Li J. Hotair facilitates hepatic stellate cells activation and fibrogenesis in the liver. Biochim Biophys Acta Mol Basis Dis 2017; 1863: 674–686.

64. Marek G, Collinsworth A, Liu C, Brantly M, Clark V. Quantitative measurement of the histological features of alpha-1 antitrypsin deficiency-associated liver disease in biopsy specimens. PLoS One 2021; 16: e0256117.

65. Zhang Y, Guo J, Li Y, Jiao K, Zhang Y. let-7a suppresses liver fibrosis via TGFbeta/SMAD signaling transduction pathway. Exp Ther Med 2019; 17: 3935–3942.

66. Gamez-Valero A, Campdelacreu J, Vilas D, Ispierto L, Rene R, Alvarez R, Armengol MP, Borras FE, Beyer K. Exploratory study on microRNA profiles from plasma-derived extracellular vesicles in Alzheimer’s disease and dementia with Lewy bodies. Transl Neurodegener 2019; 8: 31.

67. Liu J, Fan L, Yu H, Zhang J, He Y, Feng D, Wang F, Li X, Liu Q, Li Y, Guo Z, Gao B, Wei W, Wang H, Sun G. Endoplasmic Reticulum Stress Causes Liver Cancer Cells to Release Exosomal miR-23a-3p and Up-regulate Programmed Death Ligand 1 Expression in Macrophages. Hepatology 2019; 70: 241–258.

68. Sun LY, Wang N, Ban T, Sun YH, Han Y, Sun LL, Yan Y, Kang XH, Chen S, Sun LH, Zhang R, Zhao YJ, Zhang H, Ai J, Yang BF. MicroRNA-23a mediates mitochondrial compromise in estrogen deficiency-induced concentric remodeling via targeting PGC-1alpha. J Mol Cell Cardiol 2014; 75: 1–11.

69. Li L, Zhang X, Ren H, Huang X, Shen T, Tang W, Dou L, Li J. miR-23a/b-3p promotes hepatic lipid accumulation by regulating Srebp-1c and Fas. J Mol Endocrinol 2021; 68: 35–49.

70. Lu Y, Wang LR, Lee J, Mohammad NS, Aranyos AM, Gould C, Khodayari N, Oshins RA, Moneypenny CG, Brantly ML. The unfolded protein response to PI*Z alpha-1 antitrypsin in human hepatocellular and murine models. Hepatol Commun 2022; 6: 2354–2367.

71. Mitchell EL, Khan Z. Liver Disease in Alpha-1 Antitrypsin Deficiency: Current Approaches and Future Directions. Curr Pathobiol Rep 2017; 5: 243–252.

72. Khodayari N, Wang RL, Oshins R, Lu Y, Millett M, Aranyos AM, Mostofizadeh S, Scindia Y, Flagg TO, Brantly M. The Mechanism of Mitochondrial Injury in Alpha-1 Antitrypsin Deficiency Mediated Liver Disease. Int J Mol Sci 2021; 22.

73. Teckman JH, An JK, Blomenkamp K, Schmidt B, Perlmutter D. Mitochondrial autophagy and injury in the liver in alpha 1-antitrypsin deficiency. Am J Physiol Gastrointest Liver Physiol 2004; 286: G851–862.

74. Vychytilova-Faltejskova P, Radova L, Sachlova M, Kosarova Z, Slaba K, Fabian P, Grolich T, Prochazka V, Kala Z, Svoboda M, Kiss I, Vyzula R, Slaby O. Serum-based microRNA signatures in early diagnosis and prognosis prediction of colon cancer. Carcinogenesis 2016; 37: 941–950.

75. Kaserman JE, Werder RB, Wang F, Matte T, Higgins MI, Dodge M, Lindstrom-Vautrin J, Bawa P, Hinds A, Bullitt E, Caballero IS, Shi X, Gerszten RE, Brunetti-Pierri N, Liesa M, Villacorta-Martin C, Hollenberg AN, Kotton DN, Wilson AA. Human iPSC-hepatocyte modeling of alpha-1 antitrypsin heterozygosity reveals metabolic dysregulation and cellular heterogeneity. Cell Rep 2022; 41: 111775.

76. Vickers KC, Landstreet SR, Levin MG, Shoucri BM, Toth CL, Taylor RC, Palmisano BT, Tabet F, Cui HL, Rye KA, Sethupathy P, Remaley AT. MicroRNA-223 coordinates cholesterol homeostasis. Proc Natl Acad Sci U S A 2014; 111: 14518–14523.

77. Chu M, Zhao Y, Yu S, Hao Y, Zhang P, Feng Y, Zhang H, Ma D, Liu J, Cheng M, Li L, Shen W, Cao H, Li Q, Min L. miR-15b negatively correlates with lipid metabolism in mammary epithelial cells. Am J Physiol Cell Physiol 2018; 314: C43–C52.

78. Doumatey AP, He WJ, Gaye A, Lei L, Zhou J, Gibbons GH, Adeyemo A, Rotimi CN. Circulating MiR-374a-5p is a potential modulator of the inflammatory process in obesity. Sci Rep 2018; 8: 7680.

79. Lang A, Grether-Beck S, Singh M, Kuck F, Jakob S, Kefalas A, Altinoluk-Hambuchen S, Graffmann N, Schneider M, Lindecke A, Brenden H, Felsner I, Ezzahoini H, Marini A, Weinhold S, Vierkotter A, Tigges J, Schmidt S, Stuhler K, Kohrer K, Uhrberg M, Haendeler J, Krutmann J, Piekorz RP. MicroRNA-15b regulates mitochondrial ROS production and the senescence-associated secretory phenotype through sirtuin 4/SIRT4. Aging (Albany NY) 2016; 8: 484–505.

80. Nishi H, Ono K, Iwanaga Y, Horie T, Nagao K, Takemura G, Kinoshita M, Kuwabara Y, Mori RT, Hasegawa K, Kita T, Kimura T. MicroRNA-15b modulates cellular ATP levels and degenerates mitochondria via Arl2 in neonatal rat cardiac myocytes. J Biol Chem 2010; 285: 4920–4930.

81. Sun Z, Gao Z, Wu J, Zheng X, Jing S, Wang W. MSC-Derived Extracellular Vesicles Activate Mitophagy to Alleviate Renal Ischemia/Reperfusion Injury via the miR-223-3p/NLRP3 Axis. Stem Cells Int 2022; 2022: 6852661.

82. Vulf M, Shunkina D, Komar A, Bograya M, Zatolokin P, Kirienkova E, Gazatova N, Kozlov I, Litvinova L. Analysis of miRNAs Profiles in Serum of Patients With Steatosis and Steatohepatitis. Front Cell Dev Biol 2021; 9: 736677.

83. Gu C, Hou C, Zhang S. miR-425-5p improves inflammation and septic liver damage through negatively regulating the RIP1-mediated necroptosis. Inflamm Res 2020; 69: 299–308.

84. Nakagawa R, Muroyama R, Saeki C, Goto K, Kaise Y, Koike K, Nakano M, Matsubara Y, Takano K, Ito S, Saruta M, Kato N, Zeniya M. miR-425 regulates inflammatory cytokine production in CD4(+) T cells via N-Ras upregulation in primary biliary cholangitis. J Hepatol 2017; 66: 1223–1230.

85. Rao D, Guan S, Huang J, Chang Q, Duan S. miR-425-5p Acts as a Molecular Marker and Promoted Proliferation, Migration by Targeting RNF11 in Hepatocellular Carcinoma. Biomed Res Int 2020; 2020: 6530973.

86. Hiller AM, Ekstrom M, Piitulainen E, Lindberg A, Ronmark E, Tanash H. Cancer risk in severe alpha-1-antitrypsin deficiency. Eur Respir J 2022; 60.

87. Zhang CY, Yuan WG, He P, Lei JH, Wang CX. Liver fibrosis and hepatic stellate cells: Etiology, pathological hallmarks and therapeutic targets. World J Gastroenterol 2016; 22: 10512–10522.

88. Wang X, Tang X, Gong X, Albanis E, Friedman SL, Mao Z. Regulation of hepatic stellate cell activation and growth by transcription factor myocyte enhancer factor 2. Gastroenterology 2004; 127: 1174–1188.

89. Tas Hekimoglu A, Toprak G, Akkoc H, Evliyaoglu O, Ozekinci S, Kelle I. Oxytocin ameliorates remote liver injury induced by renal ischemia-reperfusion in rats. Korean J Physiol Pharmacol 2013; 17: 169–173.

90. Luo D, Jin B, Zhai X, Li J, Liu C, Guo W, Li J. Oxytocin promotes hepatic regeneration in elderly mice. iScience 2021; 24: 102125.

91. Wang Y, Lin Z, Wan H, Zhang W, Xia F, Chen Y, Chen X, Wang C, Chen C, Wang N, Lu Y. Glucagon is associated with NAFLD inflammatory progression in type 2 diabetes, not with NAFLD fibrotic progression. Eur J Gastroenterol Hepatol 2021; 33: e818–e823.

92. Suppli MP, Rigbolt KTG, Veidal SS, Heeboll S, Eriksen PL, Demant M, Bagger JI, Nielsen JC, Oro D, Thrane SW, Lund A, Strandberg C, Konig MJ, Vilsboll T, Vrang N, Thomsen KL, Gronbaek H, Jelsing J, Hansen HH, Knop FK. Hepatic transcriptome signatures in patients with varying degrees of nonalcoholic fatty liver disease compared with healthy normal-weight individuals. Am J Physiol Gastrointest Liver Physiol 2019; 316: G462–G472.

93. Tai Y, Zhao C, Lan T, Zhang L, Xiao Y, Tong H, Liu R, Tang C, Gao J. Integrated Analysis of Hepatic miRNA and mRNA Expression Profiles in the Spontaneous Reversal Process of Liver Fibrosis. Front Genet 2021; 12: 706341.

94. Aass KR, Nedal TMV, Tryggestad SS, Haukas E, Slordahl TS, Waage A, Standal T, Mjelle R. Paired miRNA-and messenger RNA-sequencing identifies novel miRNA-mRNA interactions in multiple myeloma. Sci Rep 2022; 12: 12147.

95. Han X, Wu Y, Yang Q, Cao G. Peroxisome proliferator-activated receptors in the pathogenesis and therapies of liver fibrosis. Pharmacol Ther 2021; 222: 107791.

96. Westenberger G, Sellers J, Fernando S, Junkins S, Han SM, Min K, Lawan A. Function of Mitogen-Activated Protein Kinases in Hepatic Inflammation. J Cell Signal 2021; 2: 172–180.

97. Troeger JS, Mederacke I, Gwak GY, Dapito DH, Mu X, Hsu CC, Pradere JP, Friedman RA, Schwabe RF. Deactivation of hepatic stellate cells during liver fibrosis resolution in mice. Gastroenterology 2012; 143: 1073–1083 e1022.

98. Song YM, Lee YH, Kim JW, Ham DS, Kang ES, Cha BS, Lee HC, Lee BW. Metformin alleviates hepatosteatosis by restoring SIRT1-mediated autophagy induction via an AMP-activated protein kinase-independent pathway. Autophagy 2015; 11: 46–59.

99. Reis-Barbosa PH, Marinho TS, Matsuura C, Aguila MB, de Carvalho JJ, Mandarim-de-Lacerda CA. The obesity and nonalcoholic fatty liver disease mouse model revisited: Liver oxidative stress, hepatocyte apoptosis, and proliferation. Acta Histochem 2022; 124: 151937.

100. Lambrecht J, Jan Poortmans P, Verhulst S, Reynaert H, Mannaerts I, van Grunsven LA. Circulating ECV-Associated miRNAs as Potential Clinical Biomarkers in Early Stage HBV and HCV Induced Liver Fibrosis. Front Pharmacol 2017; 8: 56.

